# Synaptic scale dopamine disruption in Huntington’s Disease model mice imaged with near infrared catecholamine nanosensors

**DOI:** 10.1101/2022.09.19.508617

**Authors:** Sarah J. Yang, Jackson Travis del Bonis O’Donnell, Francesca Giordani, Jeffery Wang, Alison Lui, David Piekarski, Ashvin Irrinki, David V. Schaffer, Markita P. Landry

## Abstract

Dopamine neuromodulation is a critical process that facilitates learning, motivation, and motor control. Disruption of these processes has been implicated in several neurological and psychiatric disorders including Huntington’s Disease (HD). While several treatments for physical and psychiatric HD symptoms target dopaminergic neuromodulation, the mechanism by which dopaminergic dysfunction occurs during HD is unknown. This is partly due to limited capability to visualize dopamine dynamics at the spatiotemporal resolution of both neuromodulator release (ms) and dopaminergic boutons (µm). Here we employ near-infrared fluorescent catecholamine nanosensors (nIRCats) to image dopamine release within the brain striatum of R6/2 Huntington’s Disease Model (R6/2) mice. We find that stimulated dorsal striatal dopamine release decreases with progressive motor degeneration and that these decreases are primarily driven by a decrease in the number of dopamine hotspots combined with decreased release intensity and decreased release fidelity. Using nIRCat’s high spatial resolution, we show that dopamine hotspots in late HD show increased ability to add new dopamine hotspots at high extracellular calcium concentrations and track individual dopamine hotspots over repeated stimulations and pharmacological wash to measure dopamine hotspots release fidelity. Compellingly, we demonstrate that antagonism of D2-autoreceptors using Sulpiride and direct blocking of K_v_1.2 channels using 4-Aminopyradine (4-AP) increases the fidelity of dopamine hotspot activity in WT striatum but not in late HD striatum, indicating that D2-autoreceptor regulation of dopamine release through K_v_1.2 channels is compromised in late HD. These findings, enabled by nIRCats, provide a more detailed look into how dopamine release is disrupted and dysregulated during Huntington’s Disease to alter the coverage of dopamine modulation across the dorsal striatum.

**SIGNIFICANCE STATEMENT:** Huntington’s Disease (HD) is a neurodegenerative disorder with no cure. Dysfunction of dopamine signaling is known to deteriorate in HD but has not been studied at the spatial level of individual release sites. Here, we image dopamine release from individual hotspots in brain slices from R6/2 HD mice at early and late disease timepoints with dopamine nanosensors. We track single dopamine hotspots and find that dopamine hotspot number, release intensity, and release fidelity decrease in HD, and demonstrate that changes in D2-autoreceptor regulation manifest through changes in hotspot release fidelity thus compromising dopamine coverage across the dorsal lateral striatum. These findings highlight dopaminergic neurons in cortico-striatal signaling during HD as a promising new therapeutic target for HD treatment.

## INTRODUCTION

Huntington’s Disease (HD) is a genetic, neurodegenerative disorder caused by aberrant expansion of the CAG (glutamine) repeat region of the Huntingtin Gene (HTT) (Finkbeiner, 2011). Patients with HD characteristically present with motor dysfunction as well as cognitive and psychiatric disorders beginning at early adulthood (ages 20-30) (Finkbeiner, 2011). Initial motor dysfunction is characterized by *chorea*, non-voluntary dance-like movements, and gradually transitions into *bradykinesia* late in disease. Neurodegeneration in HD occurs primarily in the Striatum — a brain structure critical to the relay of volitional movement — via the selective dysfunction and death of medium spiny neurons (MSN). This dysfunction is often coincident with the formation of mutant huntingtin protein (mhtt) aggregates, though whether mhtt aggregate function as protective or disease-causing agents remains unknown (Arrasate et al., 2004; Takahashi et al., 2008). Interestingly, huntingtin is ubiquitously expressed throughout the brain and mhtt does not show selective change in expression in MSNs (Li et al., 1993; DiFiglia et al., 1995; Landwehrmeyer et al., 1995). Electrophysiological studies have shown that neuronal signaling is disrupted during HD, with aberrant glutamatergic inputs from the cortex onto MSNs having potential excitotoxic effects (André et al., 2010; Rangel-Barajas and Rebec, 2016). As such, striatal degeneration and behavioral changes that manifest during HD may arise through combined cell autonomous effects and synaptic dysfunction (Cepeda and Levine, 2020).

In this light, understanding the nature of synaptic dysfunction during HD is vital to expanding our understanding of HD. Healthy striatal function relies on dopamine release from neurons projecting from the Substantia Nigra pars compacta (SNc) to potentiate glutamatergic synapses onto direct and indirect pathway MSNs via dopamine D1 Receptors (D1R) and dopamine D2 Receptors (D2Rs) (Bariselli et al., 2019). Decreases in dopamine tone and release, as in the case of Parkinson’s Disease, results in impaired motor function (Segura-Aguilar et al., 2014). Similarly, bi-phasic changes in dopamine release has been noted in both human HD patients and HD animal models, often with elevated dopamine release coinciding with choreic motor phenotypes and decreased dopamine release coinciding with bradykinesia (Ortiz et al., 2010, 2011; Callahan and Abercrombie, 2011; Cepeda et al., 2014). Though there is no present cure for HD, treatments aimed towards symptom management primarily target dopaminergic, glutamate or GABA signaling (Frank, 2014). Novel therapies principally seek to decrease the amount of mutant huntingtin protein in patients or replacing degenerated neurons (Machida et al., 2006; McBride et al., 2011; Carri et al., 2013; Fink et al., 2016; Adil et al., 2018; Evers et al., 2018; Ekman et al., 2019). However, to date, these therapeutics have yet to demonstrate efficacy in clinical trials (Kwon, 2021; Sheridan, 2021). These efforts are largely directed towards the cortex and striatum, areas of noted degeneration in HD but distal to the location of dopaminergic cell bodies in the substantia nigra pars compacta.

While general trends in dopamine levels have been reported for HD, comparatively little is known about how dopaminergic signaling changes at the level of release sites. Recent findings have shown that some portion of striatal dopamine release arises from defined axonal sites equipped with fast-release synaptic machinery (Liu et al., 2018; Banerjee et al., 2020). Simulations of dopamine release have also shown the importance of dopaminergic coverage across the striatum for effective activation of D1-Receptors (D1R) and D2-Rreceptors (D2R) on MSNs (Dreyer et al., 2010; Dreyer and Hounsgaard, 2013). Challenges in measuring dopamine release at this level of spatial resolution has historically been in part due to lack of tools for high spatio-temporal imaging. Recently, genetically encoded dopamine sensors have shown promise in imaging spatially defined dopamine release *in vivo* and *ex vivo* (Sun et al., 2018; Patriarchi et al., 2019). However, due to these sensors’ structural similarities to endogenous dopamine receptors, they are not compatible for studies incorporating pharmacological drugs directed towards dopamine receptors (dopamine pharmacology). Previously we have shown that single-walled carbon nanotube sensors such as the near infrared catecholamine sensor (nIRCats) serve as adept, dopamine pharmacology compatible sensors in the dorsal striatum capable of imaging 2 µm dopamine release sites (Beyene et al., 2019; Yang et al., 2020). The non-genetically encoded aspect of these sensors allows them to be readily deployed in disease model animals and at a wide range of ages. Here we conduct dopamine nIRCat imaging in *ex vivo* brain slices taken from 4 week, 9 week, and 12 week R6/2 HD disease model mice that are known to undergo progressive decrease in dopamine tone and release ability along with progressive motor degeneration (Johnson et al., 2006, 2007; Ortiz et al., 2010; Callahan and Abercrombie, 2011; Kaplan et al., 2018). We also explore the effect of external calcium concentration on dopamine release before and after motor symptom onset, disease-related changes in D2-autoreceptor regulation of dopamine release using D2R antagonist Sulpiride, and changes in K_v_1.2 channel function using 4-Aminopyradine (4AP).

## MATERIALS AND METHODS

### Animals

Male B6CBA-Tg(HDexon1)62Gpb/3J mice (R6/2 mice) were purchased from Jackson Labs and bred at 6 weeks with 10 week old female C57BL/6 mice. Pups were weaned and genotyped for the human HD fragment at 3 weeks. Mice were housed at three to five animals per cage with food and water available *ad libitum* and maintained in a temperature-controlled environment on a 12h dark/light cycle with light-on at 7:00 am and light-off at 7:00 pm. All animal procedures were approved by the University of California Berkeley Animal Care and Use Committee.

### nIRCat Nanosensor synthesis and characterization

Dopamine nIRCat nanosensor was synthesized and characterized as described previously described in (Yang et al., 2021). A single walled carbon nanotube (SWNT) slurry was created by combining 1050 mg of hydrated HiPco SWNTs purchased from NanoIntegris with 25 mL of molecular grade water in a 50 mL Falcon Tube and probe sonicating the solution for 2 minutes at 10% amplitude until the slurry is visually distributed. To create nIRCat nanosensors, 100 µl of SWNT slurry was mixed with 1 mg of (GT)_6_ oligonucleotides purchased from Integrated DNA Technologies (standard desalting) in 100 mM and bath sonicated for 10 minutes (Branson Ultrasonic 1800) followed by 5 minutes of rest at room temperature. The solution was then sonicated on ice for 10 minutes using a probe-tip sonicator (Cole-Parmer Ultrasonic Processor, 3-mm diameter tip, 5 W power) followed by 5 minutes of rest on ice. The sonicated solution was incubated at room temperature for 30 mins and centrifuged at 16,000 g (Eppendorf 5418) for 30 minutes to removed unsuspended SWNT bundles and amorphous carbon. The supernatant is the removed for use and stored at 4°C for 30 minutes before characterization. Final supernatant should be stored at 4°C until use.

Nanosensors are synthesized in 1 mL batches and combined for characterization. Nanosensor concentrations were determined using absorbance at 632 nM with an extinction coefficient of ε = 0.036 (mg/L)^-1^cm^-1^. To characterize the visible and nIR absorption spectrum, nanosensors were diluted to a concentration of 5 mg/L in 1x PBA and taken using a UV-VIS-nIRC spectrophotometer (Shimadzu UV-3600 Plus). To test fluorescent response to dopamine administration, each sensor batch is diluted to a working concentration of 5 mg/L in 1x PBS and 198 µl aliquots are made into a 96-well plate and baseline fluorescence is taken using a 20x objective on an inverted Zeiss microscope (Axio Observer D1) coupled to a Princeton Instruments spectrograph (SCT 320) and a liquid nitrogen cooled Princeton Instruments InCaAs linear array detector (PyLoN-IR). Nanosensors were excited using a 721-nm lazer (Opto Engine LLC). After the baseline fluorescence was taken, 2 µl of 10 mM Dopamine in 1xPBS is added and a robust fluorescence response to dopamine was confirmed.

### Phenotypic Motor Coordination Assessment

The accelerating Rotarod test and hind limb clasp test were used to evaluate changes in motor coordination in R6/2 and WT mice. For accelerating rotarod tests, mice were placed on a Ugo Basile rotarod for 1 min a 5 rpm to adjust to the apparatus. At the end of the 1 min adjustment period, the speed of the rotarod was increased at a constant rate to a final speed of 40 rpm over 350 s. The trial is terminated after mice either fall off the rod, tumble on the rod for two consecutive rotations, or “max out” the rod speed at 360s. Starting at four weeks, mice are introduced to the rotarod and complete the test for 3 consecutive days, before their rotarod times plateau and performance is recorded on the fourth day. For subsequent weeks, mice complete the rotarod only once a week.

Hind limb clasp tests are conducted by grasping mice at the base of the tail and lifting the mouse off the ground for 10 s. Mice that show splayed out legs are assigned a score of 0, mice that contract one hindlimb are scored at 1, mice contract both hindlimbs are scored at 2, and mice that retract both hindlimbs full and curl into the abdomen are scored at 3.

### nIRCat dopamine Imaging

Acute live brain slices were prepared using protocols previously described (Yang et al., 2021). Briefly, mice are deeply anesthetized via intraperitoneal ketamine/xylazine cocktail and perfused transcardially using cold cutting buffer (119 mM NaCl, 26.2 mM NaHCO3, 2.5 mM KCl, 1 mM NaH2PO4,3.5 mM MgCl2, 10 mM glucose, and 0 mM CaCl2). The brain was then rapidly dissected, mounted on a vibratome stage (Leica VT1200 S) using super glue, and cut into 300 µm thick slices containing the dorsal striatum. Slices were then collected and incubated at 37°C for 30 minutes in oxygen saturated ACSF (119 mM NaCl, 26.2 mM NaHCO3, 2.5 mM KCl, 1 mM NaH2PO4, 1.3 mM MgCl2, 10 mM glucose, and 2 mM CaCl2) followed by 30-minute incubation at room temperature. All slices are maintained at room temperature until imaging and used within 6 hours of preparation.

Slices are labeled through passive incubation in 5 ml of ACSF containing nIRCat nanosensor at a concentration of 2 mg/L for 15 minutes. After incubation, the slices is transferred through 3 wells of a 24-well plate containing ACSF to rinse off non-localized nIRCat sensor and then left to rest at room temperature ACSF for 15 minutes before transfer to the 32°C recording chamber. Once placed in the recording chamber, slices equilibrate for 15 minutes during which a tungsten bipolar stimulation electrode is positioned at a field of view in the dorsal-lateral striatum using a 4x objective (Olympus XLFluor 4/ 340). Under a 60x objective the electrode is moved 200 µm away from the selected field of view and brought into contact with the surface of the brain slice. In all experiments, 600 total images are acquired into an image-stack at a rate of 9 frames per second. A single stimulation of 0.1 mA or 0.3 mA is applied after 200 frames of baseline are collected. Videos of stimulation at each strength are collected in triplicate and stimulation strengths are alternated. All slices are given 5 minutes between each stimulation with the excitation laser path shuttered. Prior to stimulation, the laser is un-shuttered for 1 minutes.

### nIRCat Imaging Calcium Wash and Sulpiride wash

To image nIRCat-labeled acute brains slices at multiple extracellular calcium concentrations, buffers were prepared at three calcium concentrations: 1 mM Low Calcium Buffer (119 mM NaCl, 26.2 mM NaHCO3, 2.5 mM KCl, 1 mM NaH2PO4, 1.3 mM MgCl2, 10 mM glucose, and 1 mM CaCl2), 2 mM Normal Calcium Buffer (119 mM NaCl, 26.2 mM NaHCO3, 2.5 mM KCl, 1 mM NaH2PO4, 1.3 mM MgCl2, 10 mM glucose, and 2 mM CaCl2), 4 mM High Calcium Buffer (119 mM NaCl, 26.2 mM NaHCO3, 2.5 mM KCl, 1 mM NaH2PO4, 1.3 mM MgCl2, 10 mM glucose, and 4 mM CaCl2). Following stimulation in 2 mM Normal Calcium Buffer, 4 mM High Calcium buffer was flowed into the imaging chamber for 15 minutes (Full bath turnover in ∼3 minutes). After buffer transfer, the slice was stimulated at 0.1 mA and 0.3 mA in triplicate as described for 2 mM Normal Calcium Buffer. Buffer was then exchanged again to 1 mM Low Calcium Buffer via 15-minute wash and the slice was stimulated at 0.1 mA and 0.3 mA in triplicate.

To nIRCat image acute brain slices in the presence of the D2-antagonist Sulpiride, S-Sulpiride was dissolved in sterile DMSO and frozen in 100 µl aliquots at −20°C. Prior to use, single aliquots are thawed and added to 100 mL of ACSF to produce a 10 µM Sulpiride solution. Acute brain slices were stimulated at 0.1 mA and 0.3 mA in triplicate in sulpiride-free ACSF. Sulpiride solution was flowed into the imagine chamber for 15 minutes before stimulating the slice at 0.1 mA and 0.3 mA in triplicate.

### Image Stack Processing and Data Analysis of nIRCat Data

Raw Image stack files are processed using a custom-built, publicly available MATLAB program (https://github.com/jtdbod/Nanosensor-Imaging-App). Image processing procedures are described in depth in Yang, del Bonis O’Donnel et al and briefly summarized here. Regions of dopamine release are identified by large changes in nIRCat ΔF/F response. To minimize bias and improve stack processing time, regions of high ΔF/F response (dopamine hotspots) were identified by defining a grid of 2 µm squares across the field of view. For each grid square ΔF/F was calculated using the formula (F-F_0_) /F_0_, where F_0_ is defined by the average fluorescence of the grid square over the first 30 frames of the image stack and F is the fluorescence intensity of the gird square as it changes over the 600 collected frames. Grid squares are identified as regions of interest if they exhibit behavior that is 3 standard deviations above the baseline F_0_ activity around time of stimulation (200 frames).

Dopamine hotspots were identified for each stimulation replicate image stack taken at a given field-of-view on a brain slice. The peak ΔF/F of each dopamine hotspot in the image stack were averaged to give the average image stack peak ΔF/F. The average image stack peak ΔF/F from the three stimulation replicates were then average to give the slice average peak ΔF/F. Similarly, the number of dopamine hotspots identified from each stimulation replicate image stack were averaged to give the slice average hotspot number. Mean dopamine release and reuptake traces are produced by averaging the average traces from each slice (3 stimulations per slice, 1 slice per animal). Percent change in hotspots was calculated as (# hotspots wash - # hotspots 2 mM Ca^+2^)/ (# hotspots 2 mM Ca^+2^), whereas change in hotspots number was calculated as (# hotspots wash - # hotspots 2 mM Ca^+2^).

To track hotspot fidelity, each initially defined grid square was assigned a unique position number, allowing the position of each identified dopamine hotspot within an image stack to be recorded. For a set of triplicate image stacks, an array of all unique hotspots active across the stimulation replicates was generated. Then python code was used to analyze whether each unique hotspot was active in each stimulation replicate. The number of stimulations a unique hotspot was active in was summed across the three replicates and assigned as the dopamine release fidelity (e.g. hotspot ‘12’ is active in 2 out of 3 stimulations and is assigned release fidelity 2). The same procedure was used to identify the dopamine release fidelity of hotspots active after drug wash. Hotspots were then separated into three groups: hotspots that are active both before and after drug wash (shared hotspots), hotspots that become active after drug wash (added hotspots), and hotspots that are only active before drug wash. For shared hotspots modulation in hotspot release strength was calculated as the difference in peak ΔF/F of the unique hotspot before and after drug wash, (mean ΔF/F)_post_ - (mean ΔF/F)_pre,_ where (mean ΔF/F)_pre_ is the average peak ΔF/F of each unique dopamine hotspot across the three stimulations before drug wash and (mean ΔF/F)_post_ is the average peak ΔF/F of each unique dopamine hotspot across the three stimulations after drug wash. For hotspots active only after drug wash, there is no corresponding “pre drug wash” ΔF/F. Therefore, the difference in peak ΔF/F was calculated through (mean ΔF/F)_post_ - (mean ΔF/F)_pre, shared_, where (mean ΔF/F)_post_ represents the average peak ΔF/F of the unique dopamine hotspot active after sulpiride wash across three stimulations and (mean ΔF/F)_pre, shared_ is the average of all the shared hotspots’ mean ΔF/F from the slice before drug wash.

#### EXPERIMENTAL DESIGN AND STATISTICAL ANALYSIS

All nIRCat Imaging data were processed using a custom-built, publicly available MATLAB program (https://github.com/jtdbod/Nanosensor-Imaging-App). Statistical analyses were conducted using the open-source statistical python package pingouin. All bar graphs show the mean with error bars denoting the 95% confidence interval. All single data points correspond to a single slice taken from an animal. Data comparing two variables was analyzed using a mixed-ANOVA with wash condition as the within-subject factor (e.g.sulpiride, blank, calcium concentration) and disease state as the between-subject factor (eg. HD, WT). Paired t-tests were used a post-hoc tests if mixed-ANOVA analyses indicated significant differences. Data comparing two values of one variable were analyzed using tukey’s t-test. Group sizes were determined based on previous literature (Adil et al., 2018). Changes in histogram skew were computed through pooling of all hotspots identified across all mice within the disease and wash condition and evaluated using a permutation test using the test statistic µ = skew(post wash) – skew(pre-wash).

#### CODE ACCESSIBILITY

All analyses were performed using in-house developed code usigin either MATLAB or python. Code to process nIRCat image stacks is available on GitHub: https://github.com/jtdbod/Nanosensor-Imaging-App.

## RESULTS

### R6/2 HD mice show progressive decrease in dopamine hotspots over disease progression and decreased individual hotspot response in late disease

Disease-related changes in dopamine signaling have been well documented in both human HD patients and multiple murine models that express mutant huntingtin protein via different avenues (Cepeda et al., 2014; Cepeda and Levine, 2020). In this work we use nIRCat imaging to investigate dopamine release in R6/2 HD mice, which have been shown through Fast Scan Cyclic Voltammetry and *in vivo* microdialysis to display decreases dopamine tone and release along with progressive decreases in motor ability typical of juvenile forms of HD (Johnson et al., 2006, 2007; Ortiz et al., 2010; Callahan and Abercrombie, 2011). We performed nIRCat dopamine imaging in R6/2 HD mice and their WT littermates at three time points: immediately at the onset of motor degeneration (p32-35, 4 wks), mid-degeneration (p64-66, 9 wk), and late in disease (p87-93, 12 wk). In parallel, we assessed the extent of motor degeneration by weekly rotarod tests, and mice were subject to nIRCat imaging at designated timepoints (Fig. 1a). Acute brain slices were incubated in the (GT)_6_ nIRCat nanosensor and subjected to single intrastriatal electrical stimulations, taken in triplicate, at both 0.3 mA and 0.1 mA. Data collected from each acute brain slice was then processed using the Neuronal Imaging Application (NIA) (Fig. 1b) (Yang et al., 2021). We have previously shown that nIRCat dopamine imaging in the dorsal striatum of acute slices reveals approximately 2µm-wide regions dopamine release hotspots, identified by sharp changes in ΔF/F fluorescence (Beyene et al., 2019). Recent studies using similar dopamine nanosensors within 2-D films have shown that these dopamine hotspots emerge from tyrosine hydroxylase (TH) positive axonal varicosities and co-localize with the pre-synaptic scaffolding protein Bassoon (Elizarova et al., 2021; Bulumulla et al., 2022). As such, we identified the average number of dopamine hotspots active within a slice during stimulation (slice average hotspot number) as well as the peak amount of dopamine released from the average hotspot within the slice (slice average peak ΔF/F) as key metrics in characterizing dopamine release dynamics (Fig. 1b).

**Figure 1.**
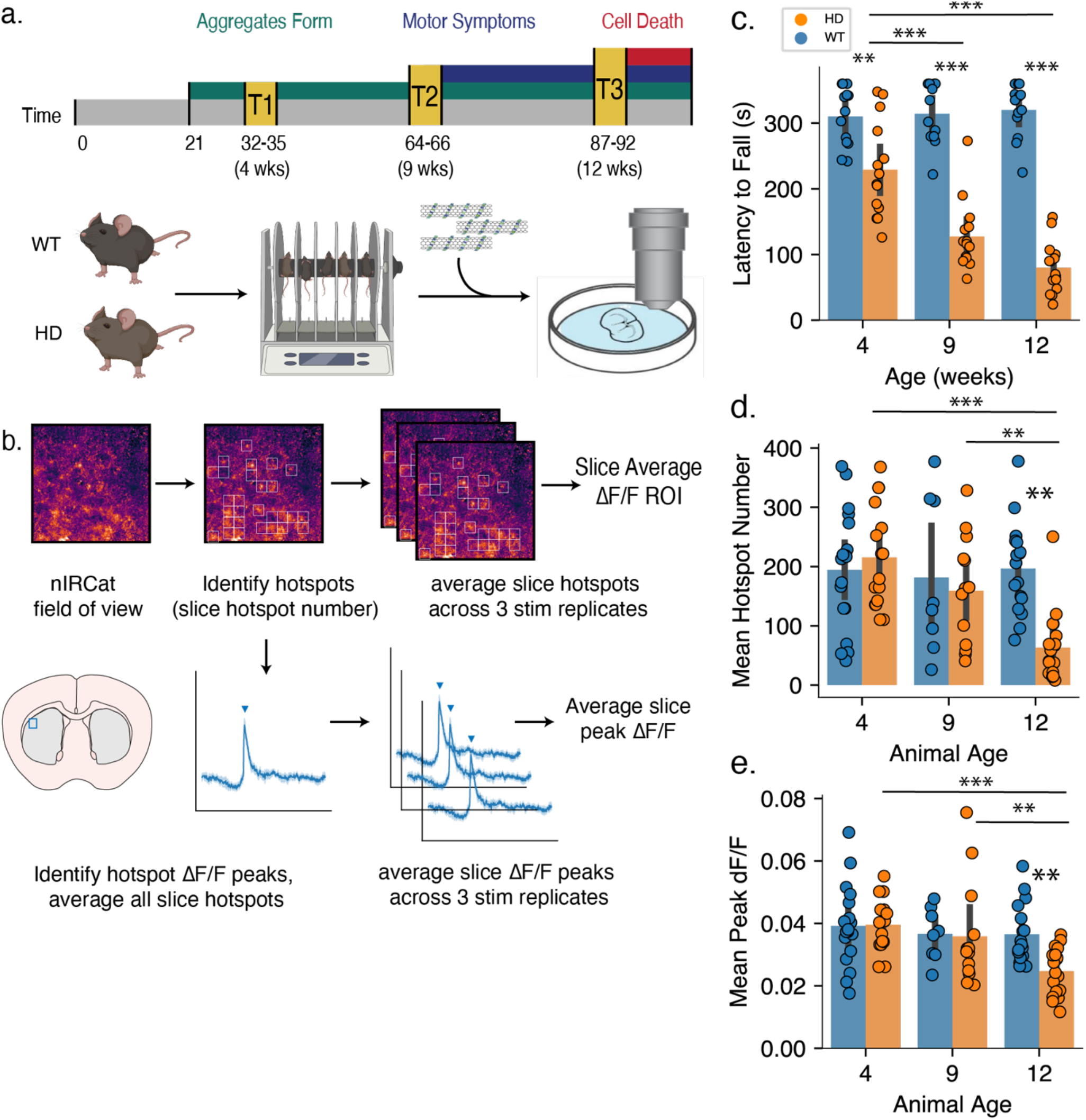
R6/2 HD mice show progressive decrease in number of dopamine hotspots over disease progression but not a change in individual dopamine ΔF/F hotspot response. *A,* Graphical overview of experimental design whereby 4 week, 9 week, and 12 week WT and R6/2 HD mice undergo weekly rotarod phenotypic assessment of motor ability followed by nIRCat dopamine imaging at the final timepoint. *B,* Graphical overview of data analysis to examine the number of putative dopamine release sites active after stimulation, termed dopamine hotspots, and the average amount of dopamine released from each site, termed average peak dopamine ΔF/F *C,* R6/2 HD mice show progressive decrease in latency to fall during an accelerating rotarod behavioral task (WT N = 13 animals, HD N = 14 animals; ANOVA: disease state, p =< 0.0005 age, p =< 0.0005; interaction, p =< 0.0005; pairwise t-test: <u>***</u> p =< 0.0005 4 wk HD/ 12 wk HD, <u>***</u> p =< 9 wk HD/12 wk HD, ns p = 0.8105 and p = 0.7531 4 wk WT/ 12 wk WT and 9 wk WT/ 12 wk WT; ** p = 0.0020 4 wk HD/4 wk WT; *** p < 0.0005 9 wk HD/9 wk WT; *** p < 0.0005 12 wk HD/12 wk WT) *D,* R6/2 HD mice show progressively decreasing numbers of dopamine hotspots from 4 weeks through 9 and 12 weeks while WT mice show no changes in dopamine hotspot number with age. (4 weeks WT N = 18 animals, HD N = 18 animals; 9 weeks WT N = 10 animals, HD N = 13 animals; 12 weeks WT N = 18 animals, HD N = 18 animals; ANOVA: disease state, p = 0.0101; animal age, p = 0.0034; interaction, p = 0.0018; pairwise t-test: <u>***</u> p < 0.0005 12wk/HD compared to 4wk/HD, <u>**</u> p =0.0037 12wk/HD compared to 9wk/HD, <u>*</u> p < 0.0005 12wk/HD compared to 12wk/WT). *E,* R6/2 HD mice show no change in average peak ΔF/F at 4 and 9 weeks but show significant decrease late in disease at 12 weeks. (ANOVA: disease state, p = 0.0469; animal age, p = 0.0047; interaction, p = 0.0530; pairwise t-test: <u>***</u> p < 0.0005 12wk/HD compared to 4wk/HD, <u>***</u> p < 0.0005 12wk/HD compared to 12wk/WT).

Consistent with findings from previous studies, WT mice showed consistent, robust performance on the rotarod across timepoints from 4 to 12 weeks while HD mice showed decreased latency to fall compared to their WT counterparts as early as 4 weeks that grew more pronounced with disease progression through 9 and 12 weeks (Fig. 1c). Dopamine release imaging with nIRCat shows that this decreasing motor ability is mirrored by decreases in the mean dopamine hotspot number and mean peak ΔF/F in HD mice (Fig. 1d, 1e). Early in disease at 4 weeks, stimulation at 0.3 mA activates a comparable number of dopamine hotspots in HD and WT mice (Fig. 1d). The dopamine hotspots observed in 4 week HD and WT mice also show similar mean peak ΔF/F, suggesting comparable dopamine release profiles (Fig. 1e). Together, these findings suggest that disruptions in rotarod performances seen at 4 weeks are not primarily driven by decreases in available dopamine. Instead, early changes in HD dorsal lateral striatal dopamine signaling may be driven by disruptions in dopamine mobilization or regulation.

In contrast, HD mice late in disease at 12 weeks show significantly fewer dopamine hotspots activated in response to 0.3 mA stimulation and decreased mean peak ΔF/F (Fig. 1d, 1e). This observed decrease in dopamine release capacity is consistent with previously established trends in R6/2 HD mice (Johnson et al., 2006, 2006, 2007; Ortiz et al., 2010; Callahan and Abercrombie, 2011). However, nIRCat’s increased spatial resolution allows new insights into the way dopamine release is compromised. These results indicate that decreased dopamine in the late R6/2 HD disease state is driven by a combination of both dopamine hotspot loss and dopamine hotspot dysfunction. This motivates exploration of molecular mechanisms implicated in dopamine hotspot activation and dopamine release as potential drivers of neurodegeneration in HD.

We also measured dopamine release at 4, 10, and 12 week timepoints at a lower intensity of 0.1 mA and found that while lower stimulation resulted in lower numbers of activated dopamine hotspots, general trends were maintained (Fig. S1a, S1b). While HD mice show trending decreases in dopamine hotspot number and slice average peak ΔF/F through 4 week to 9 weeks, there was no significant difference in either slice average hotspot number or slice average peak ΔF/F at this middle timepoint. This finding suggests that changes in other parameters of dopamine release outside those examined here may drive dysfunction at mid-disease time points.

### HD Animals at 4 weeks show increased extracellular calcium sensitivity

We next sought to examine whether the extracellular calcium sensitivity of dopamine hotspots differs between HD and WT mice early and late in disease. Changes in neuronal calcium handling have been reported in murine models of HD, though these studies have largely focused on aberrant N-methyl-D-aspartate receptor (NMDAR) signaling or mitochondrial Ca^+2^ uptake (Mackay et al., 2018). Calcium also plays a prominent role in neurotransmitter release, with release occurring when the entry of Ca^+2^ ions into the axon terminal triggers fusion of synaptic vesicles. Increased extracellular calcium concentration has been shown to modulate neurotransmitter release by increasing the probability of vesicle release, increasing the effective size of readily releasable pool vesicles, and recruiting boutons with low release probability to more active states (Leitz and Kavalali, 2011; Thanawala and Regehr, 2013). Furthermore, recent work has shown that fast, synchronous dopamine release plays a significant role in striatal dopamine signaling and that this release is mediated by the fast calcium sensor synaptotagmin-1 (Liu et al., 2018; Banerjee et al., 2020).

To examine if the calcium sensing and release machinery of dopamine hotspots is compromised during early HD, we imaged stimulated dopamine release from nIRCat labeled HD and WT slices from 4 week animals at 4 mM Ca^+2^, 2 mM Ca^+2^, and 1 mM Ca^+2^. We then examined the resulting changes in dopamine hotspot number and peak dopamine ΔF/F. Increasing extracellular Ca^+2^ concentration from 2 mM Ca^+2^ to 4 mM Ca^+2^ results in increased number of dopamine hotspots and while corresponding decrease to 1 mM Ca^+2^ results in fewer dopamine hotspots (Fig. 2a). This finding is in line with observations of glutamate release in hippocampal neurons made using pHluorin-tagged vesicles which showed that increasing extracellular Ca^+2^ concentration recruits previously low activity boutons to active dopamine hotspots (Leitz and Kavalali, 2011). We show that similar mechanisms may be involved in dorsal lateral striatal dopamine release, and that this activity can be detected by nIRCat imaging. We do not find a significant difference in the number of dopamine hotspots in HD and WT slices at any extracellular calcium concentration at 4 weeks, and both HD and WT slices at 4 weeks show a robust response to changing extracellular calcium concentration and comparable percent increase and decrease in dopamine hotspot number with increasing or decreasing extracellular Ca^+2^ concentration (Fig. 2b).

**Figure 2.**
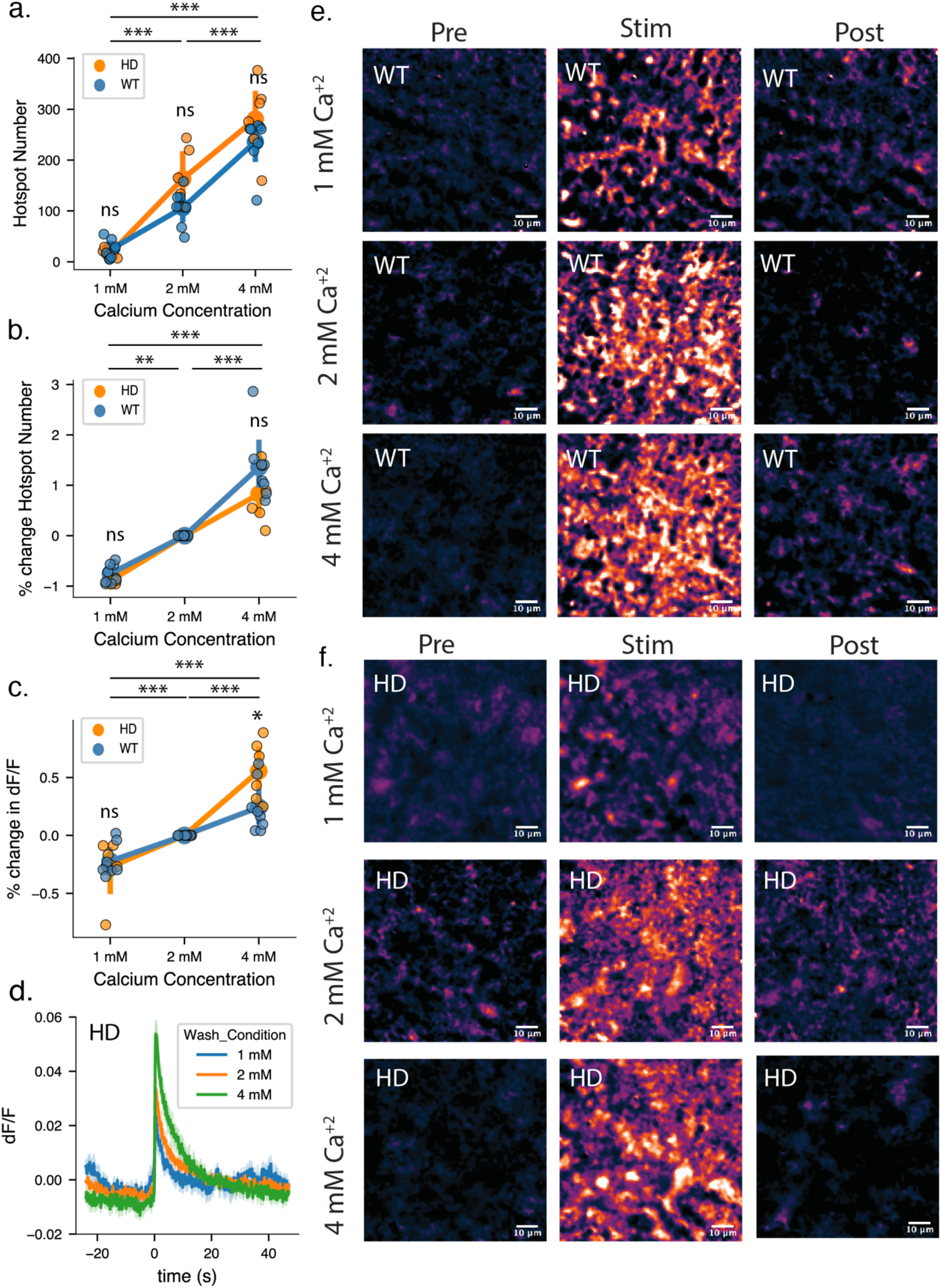
WT and R6/2 HD mice show similar extracellular calcium sensitivity for dopamine release at 4 weeks. ***A,*** The average number of dopamine hotspots active in 4 week WT and R6/2 HD striatal brain slices in response to 0.3 mA stimulation is comparable at 1 mM Ca^+2^, 2 mM Ca^+2^ and 4 mM Ca^+2^ (WT N = 9 slices, 6 animals, HD N = 6 slices, 6 animals; mixed-ANOVA: disease state, p = 0.10715; wash condition, p < 0.0005; interaction, p = 0.0735; pairwise t-test: <u>***</u> p < 0.0005 4mM Ca^+2^ compared with 1 mM Ca^+2^, <u>***</u> p < 0.0005 4mM Ca^+2^ compared with 2 mM Ca^+2^, <u>***</u> p < 0.0005 2 mM Ca^+2^ compared with 1mM Ca^+2^). ***B***, The precent change in dopamine hotspots is also comparable at all calcium concentrations. (WT N = 9 slices, 6 animals, HD N = 6 slices, 6 animals; mixed-ANOVA: disease state, p = 0.0995; wash condition, p < 0.0005; interaction, p = 0.1592; pairwise t-test: <u>***</u> p < 0.0005 4 mM Ca^+2^ compared with 1 mM Ca^+2^, <u>***</u> p < 0.0005 4 mM Ca^+2^ compared with 1 mM Ca^+2^, <u>***</u> p < 0.0005 Normal Ca^+2^ compared with Low Ca^+2^). ***B***, The percent change in dopamine hotspots is also comparable at all calcium concentrations. (WT N = 9 slices, 6 animals, HD N = 6 slices, 6 animals; mixed-ANOVA: disease state, p = 0.0995; wash condition, p < 0.0005; interaction, p = 0.1592; pairwise t-test: <u>***</u> p < 0.0005 4 mM Ca^+2^ compared with 1 mM Ca^+2^, <u>***</u> p < 0.0005 4 mM Ca^+2^ compared with 2 mM Ca^+2^, <u>***</u> p < 0.0005 2 mM Ca^+2^ compared with 1 mM Ca^+2^). ***C***, The percent increase in mean peak ΔF/F is comparable between WT and R6/2 HD striatal brain slice at 1 mM Ca^+2^ and 2 mM Ca^+2^. At 4 mM Ca^+2^ R6/2 HD slices show a 31.3% elevated response compared to WT slices (mixed-ANOVA: disease state, p = 0.2468; wash condition, p < 0.0005; interaction, p = 0.0057; pairwise t-test: <u>***</u> p < 0.0005 4 mM Ca^+2^ compared with 1 mM Ca^+2^, <u>***</u> p = 0.0002 4 mM Ca^+2^ compared with 2 mM Ca^+2^, <u>**</u> p = 0.0033 2 mM Ca^+2^ compared with 1mM Ca^+2^_;_ *p = 0.0070 High Ca^+2^/HD compared with High Ca^+2^/WT). ***D,*** Dopamine release and reuptake traces from imaged nIRCat-labeled brain slices for 4 week HD mice. Solid lines denote the average taken from all slices and light shaded bands represent one standard deviation from average behavior. A 1 ms, 0.3 mA stimulation is delivered at time = 0s. ***E,*** Representative images of dopamine release imaged in 4 week WT mice before, during, and after stimulated dopamine release. ***F,*** Representative images of dopamine release imaged in 4 week HD mice before, during, and after stimulated dopamine release.

Slice average peak dopamine ΔF/F also increases with extracellular Ca^+2^ concentration, supporting findings that high calcium concentrations increase the probability of multivesicular release events (Leitz and Kavalali, 2011). However, while 4 week HD mice do show comparable slice average peak dopamine ΔF/F to 4 week WT mice at 1 mM Ca^+2^ to 2 mM Ca^+2^, 4 week HD mice show significantly higher slice average peak dopamine ΔF/F at 4 mM Ca^+2^ (Fig. 2c, Fig. 2d). This increased calcium sensitivity in pre-symptomatic 4 week-old HD mice may suggest potential changes in calcium machinery early in HD progression that may underlie early observed changes in rotarod performance or contribute to dysfunction later in disease.

Given that increasing Ca^+2^ concentration increases peak dopamine ΔF/F, the observed increase in dopamine hotspots number at 4 mM Ca^+2^ could be driven by low releasing dopamine hotspots at 2 mM Ca^+2^ entering nIRCat’s limit of detection at 4 mM Ca^+2^. To examine this, we pooled all hotspots detected in 4 week HD slices and 4 week WT slices and plotted histograms of hotspots peak ΔF/F. Histograms for both HD and WT dopamine hotspots show a normal distribution at all calcium concentrations, suggesting that the new hotspots observed at 4 mM Ca^+2^ are not the result of low releasing hotspots entering the nIRCat’s limit of detection (Fig. S2a, Fig S2b).

### Increasing extracellular calcium in 12 wk mice increases the number of dopamine hotspots, but not to WT levels

We next investigated whether the extracellular calcium sensitivity of dopamine hotspots changes late in HD disease development. At 12 weeks, HD mice produce significantly fewer dopamine hotspots than WT mice at 2 mM Ca^+2^ and 4 mM Ca^+2^ (Fig. 3a). At 1mM Ca^+2^, low extracellular calcium concentrations are known to suppress the release probability of both HD and WT dopamine hotspots close to nIRCat’s detection limit resulting in comparable dopamine hotspot numbers (Leitz and Kavalali, 2011; Beyene et al., 2019). While increasing extracellular calcium concentration does result in an increase in dopamine hotspot number in 12 week HD slices, this increase is not sufficient to match WT levels on 2 mM Ca^+2^ or 4 mM Ca^+2^ (Fig. 3a). Interestingly, though HD slices show lower dopamine hotspot numbers than WT slices, they show a larger percent increase in hotspot number after 4 mM Ca^+2^ wash (Fig. 3b). This dopamine selective hotspot increase appears to be driven by the fact that HD and WT slices add comparable amounts of dopamine hotspots after 4 mM Ca^+2^ despite the significantly lower number of dopamine hotspots initially present in HD slices at 2 mM Ca^+2^ (Fig. S3f). In contrast to dopamine hotspot number, HD and WT slices show comparable slice average peak dopamine ΔF/F at 2 mM Ca^+2^ and decrease from WT levels at 4 mM Ca^+2^ (Fig. 3c).

**Figure 3.**
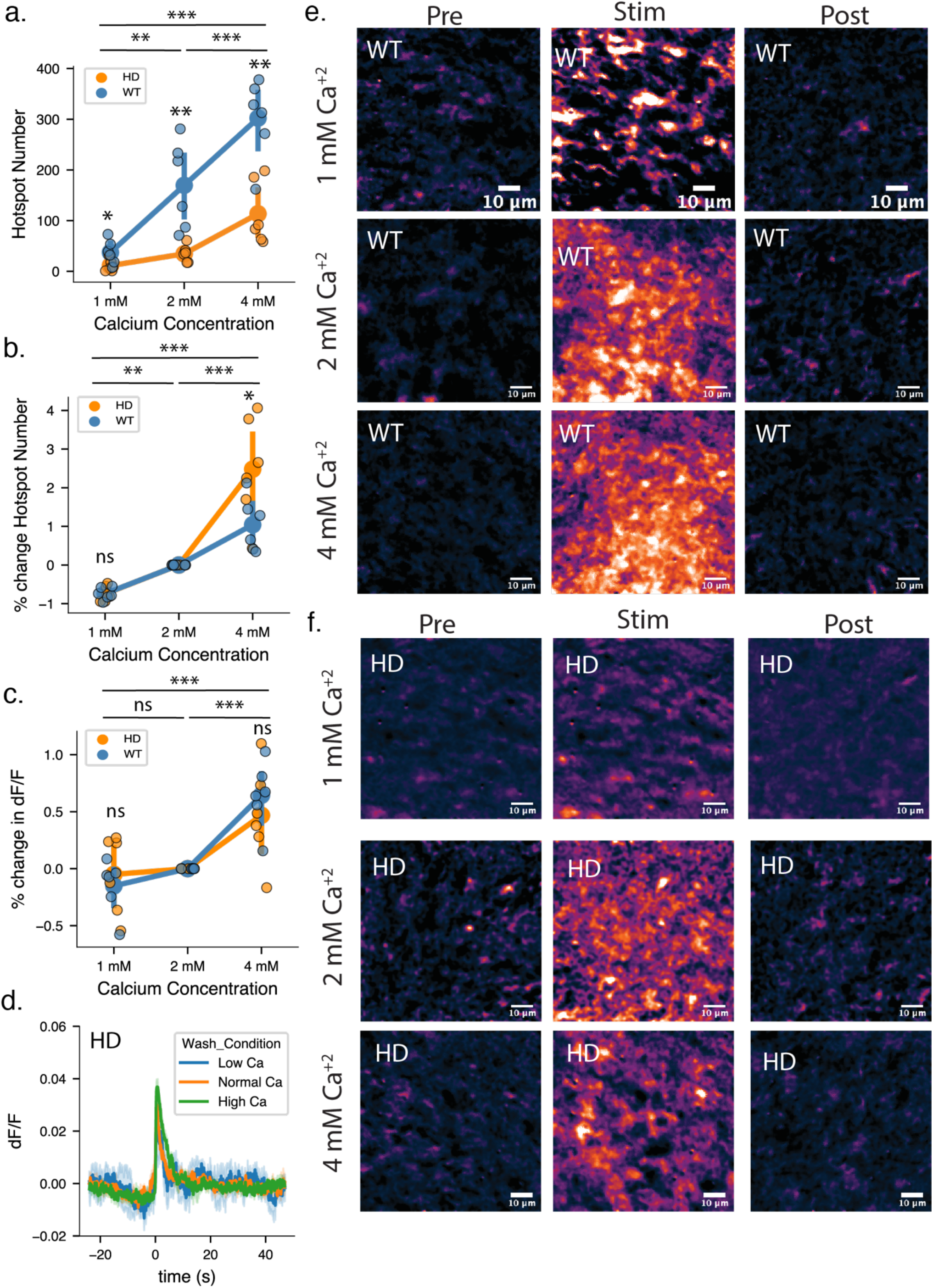
R6/2 HD mice show diminished dopamine release at 12 weeks that is improved but not fully rescued by high extracellular calcium concentration. ***A,*** The average number of dopamine hotspots active in 12 week R6/2 HD striatal brain slices in response to 0.3 mA stimulation is significantly diminished in comparison to WT brain slices. R6/2 HD slices show a 79.6% decrease in the number of dopamine hotspots at Normal Ca^+2^and a 62.4% decrease in the number of dopamine hotspots at High Ca^+2^. Increasing external calcium concentration results in an increased number of dopamine hotspots active in HD mice, but is not sufficient to fully rescue to WT levels. (WT N = 6 slices, 6 animals, HD N = 6 slices, 6 animals; mixed-ANOVA: disease state, p = 0.0010; wash condition, <u>***</u> p < 0.0005; interaction, p < 0.0005; pairwise t-test: ** p = 0.0009 HD/4 mM Ca^+2^ compared to WT/ mM Ca^+2^, ** p = 0.0036 HD/2 mM Ca^+2^ compared to WT/2 mM Ca^+2^, * p = 0.0376 HD/1 mM Ca^+2^ compared to WT/1 mM Ca^+2^). ***B,*** R6/2 HD slices show a 247.9% increase in dopamine hotspots number after 4 mM Ca^+2^ wash compared to R6/2 WT slices which show a 104.2% increase in dopamine hotspots after 4 mM Ca^+2^. (mixed-ANOVA: disease state, p = 0.0383; wash condition, <u>***</u> p < 0.0005; interaction, p = 0.0170; pairwise t-test: ** p = 0.0429 HD/High Ca^+2^ compared to WT/High Ca^+2^, <u>nr</u> p = 0.9681 HD/Low Ca^+2^ compared to WT/Low Ca^+2^). ***C,*** R6/2 HD and WT slices show comparable increase in mean peak ΔF/F at all calcium concentrations. (mixed-ANOVA: disease state, p = 0.823; wash condition, <u>***</u> p < 0.0005; interaction, p = 0.381; pairwise t-test: nr p = 0.423 HD/4 mM Ca^+2^ compared to WT/4 mM Ca^+2^, nr p = 0.568 HD/1 mM Ca^+2^ compared to WT/1 mM Ca^+2^).***D,*** Dopamine release and reuptake traces from imaged from 12 wk HD mice. Solid lines denote the average taken from all slices and light shaded bands represent one standard deviation from average behavior. A 1 ms, 0.3 mA stimulation is delivered at time = 0s. ***E,*** Representative images of dopamine release imaged in 12 week WT mice before, during, and after stimulated dopamine release. ***F,*** Representative images of dopamine release imaged in 12 week HD mice before, during, and after stimulated dopamine release.

These findings build upon existing FSCV measurements in R6/2 mice which previously reported that 12 week HD and WT mice show comparable changes in peak dopamine release concentration in response to increasing extracellular concentration (Johnson et al. 2007). The spatial insights afforded by nIRCat imaging show that while there is significant degeneration in the number of dopamine hotspots in 12 week HD slices there remains a population of dopamine hotspots in HD slices that can be made active through increasing the calcium influx into dopaminergic release sites. Furthermore, these 12 week HD slices show increased sensitivity to extracellular calcium concentration when adding new dopamine hotspots. As such, changes in calcium dependent dopamine release in late HD may play a larger role in shaping late disease states than previously expected.

### R6/2 HD mice show changes in modulation of dopamine release by D2-autoreceptor antagonist Sulpiride at 4 weeks

Axonal dopamine release in the striatum is regulated at multiple stages of the dopamine release process. As such, the amount of axonal dopamine release does not linearly scale with neuron intracellular Ca^+2^ levels (Liu and Kaeser, 2019). Striatal dopamine release can shape future release through presynaptic feedback inhibition via Dopamine Type 2 Receptors located on dopamine axons termed D2-autoreceptors (Westerink and de Vries, 1989; Sesack et al., 1994; Sulzer et al., 2016). Though the exact mechanism that underlies this feedback inhibition is still unknown, it is hypothesized that D2-autoreceptors allow for transient inhibition of dopamine release during prolonged activity through pathways involving of voltage gated calcium channels, 4-amino-pyradine (4-AP) sensitive G-protein activated inwardly rectifying potassium (GIRK) channels, modulation of dopamine synthesis by tyrosine hydroxylase (TH), and regulation of the expression of neuronal vesicular monoamine transporter (VMAT2) (Benoit-Marand et al., 2001; Schmitz et al., 2003; Sulzer et al., 2016). While decreases in broad striatal D2-receptor transcription and expression have been documented in R6/2 mice and in the caudate of human patients, less is known about how D2-autoreceptors are affected by disease course (Vashishtha et al., 2013; Achour et al., 2015). To this end, we utilize nIRCats’ compatibility with dopamine receptor pharmacology to examine D2-autoreceptor behavior using Sulpiride, a selective D2-receptor antagonist. Sulpiride is used in the treatment of Huntington’s Disease as well as Schizophrenia. However, Sulpiride’s precise mechanism of therapeutic action is presently not fully understood. Within brain slices, Sulpiride antagonism of D2-autoreceptors allows for disinhibition of dopamine synthesis and release through multiple pathways, allowing for increased dopamine release upon stimulation.

We first examined the effect of Sulpiride on HD and WT mice at 4 weeks. While the number of dopamine hotspots in HD and WT slices is not significantly different at 4 weeks, HD and WT slices do show differences in Sulpiride response (Fig. 4a). Initially, wash on of Sulpiride did not initially appear to drive an increase in the average number of dopamine hotspots in HD and WT slices wash (Fig. 4a). However, examination of the percent change in dopamine hotspot number within individual slices rather than the average number of dopamine hotspots across all slices show that Sulpiride wash drives a significant percent increase in dopamine hotspots in both HD and WT slices at 4 weeks (Fig. 4b). Intriguingly, HD slices show a larger percent increase in dopamine hotspots following Sulpiride wash than their WT counterparts. This may be in part due to observed decreases in dopamine hotspot number in a WT slices with large numbers of dopamine hotspots active in Blank ACSF. Both 4 week HD and WT slices show a comparable percent increase in mean peak ΔF/F after Sulpiride wash (Fig. 4b, Fig. 4c). Collectively, these findings suggest that changes in D2-autoreceptor expression and signaling may begin in HD slices as early as 4 weeks.

**Figure 4.**
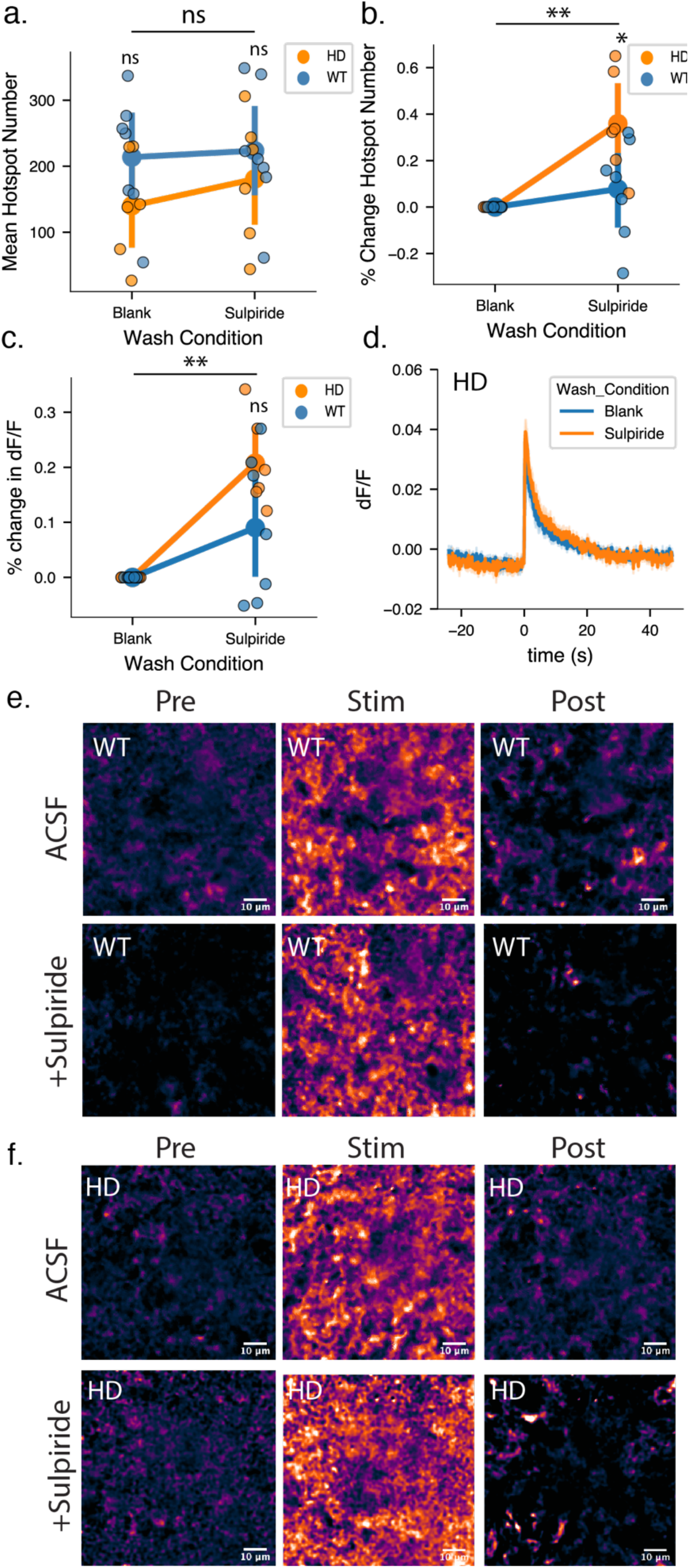
Both WT and R6/2 HD mice at 4 weeks show modulation of dopamine release via D2-autoreceptor antagonist Sulpiride. ***A,*** Both WT and R6/2 HD slices show a comparable increase in active dopamine hotspots in response to 0.3 mA stimulation after Sulpiride wash. (WT N = 7 slices, 7 animals, HD N = 6 slices, 6 animals; mixed-ANOVA: disease state, p = 0.2728; wash condition, p = 0.0733; interaction, p = 0.2313; paired t-test: nr p = 0.1589 HD/Blank compared to WT/Blank, nr p = 0.4469 HD/Sulpiride compared to WT/Sulpiride) ***B,*** R6/2 HD slices show a larger percent increase in dopamine hotspots after Sulpiride wash compared to WT slices at 4 weeks (mixed-ANOVA: disease state, p = 0.0419; wash condition, <u>**</u> p < 0.0059; interaction, p = 0.0419; paired t-test: * p = 0.0433 HD/Sulpiride compared to WT/Sulpiride). ***C,*** Both R6/2 HD and WT slices show similar increase in percent increase in peak ΔF/F after Sulpiride wash (mixed-ANOVA: disease state, p = 0.088; wash condition, p = 0.001; interaction, p = 0.0878; paired t-test: ns p = 0.080 HD/Sulpiride compared to WT/Sulpiride). ***D,*** Dopamine release and reuptake traces from 12 wk HD mice. Solid lines denote the average taken from all slices and light shaded bands represent one standard deviation from average behavior. A 1 ms, 0.3 mA stimulation is delivered at time = 0s. ***E,*** Representative images of dopamine release imaged in 4 week WT mice before, during, and after stimulated dopamine release in the presence and absence of Sulpiride. ***F,*** Representative images of dopamine release imaged in 4 week HD mice before, during, and after stimulated dopamine release in the presence and absence of Sulpiride.

### 12-week R6/2 HD mice show comparable modulation of dopamine release by D2-autoreceptor antagonist Sulpiride

We next sought to assess the Sulpiride response of HD and WT slices after advanced neurodegeneration at 12 weeks. Both HD and WT mice show increased dopamine hotspot number and slice average peak ΔF/F in response to Sulpiride wash on at 12 weeks (Fig. 5a, Fig. 5c). Interestingly, despite HD slices showing fewer active dopamine hotspots than their WT counterparts, both HD and WT slice show similar percent increase in dopamine hotspots after Sulpiride wash (Fig 5b). Furthermore, HD and WT slices show similar modulation of slice average peak dopamine ΔF/F in response to Sulpiride wash (Fig. 5b, Fig 5d). These results suggest that despite disruptions in HD D2-autoreceptor activity at 4 weeks and loss of active dopamine hotspots in HD slices at 12 weeks, D2-autoreceptor action on dopamine hotspot addition and dopamine hotspot performance of the remaining hotspots is similar between HD and WT slices late in disease. It is possible that individual pathways of D2-autoreceptor action may be differentially affected in late HD. Comparable modulation of slice average peak ΔF/F may indicate that pathways involved TH and VMAT2 which contribute to the size of dopamine release events may be relatively unaffected in late HD. Whereas disruption in mechanisms that recruit voltage gated calcium currents or 4-amino-pyradine (4-AP) sensitive K^+1^ channels in late HD may underlie decreased numbers of active dopamine hotspots.

**Figure 5.**
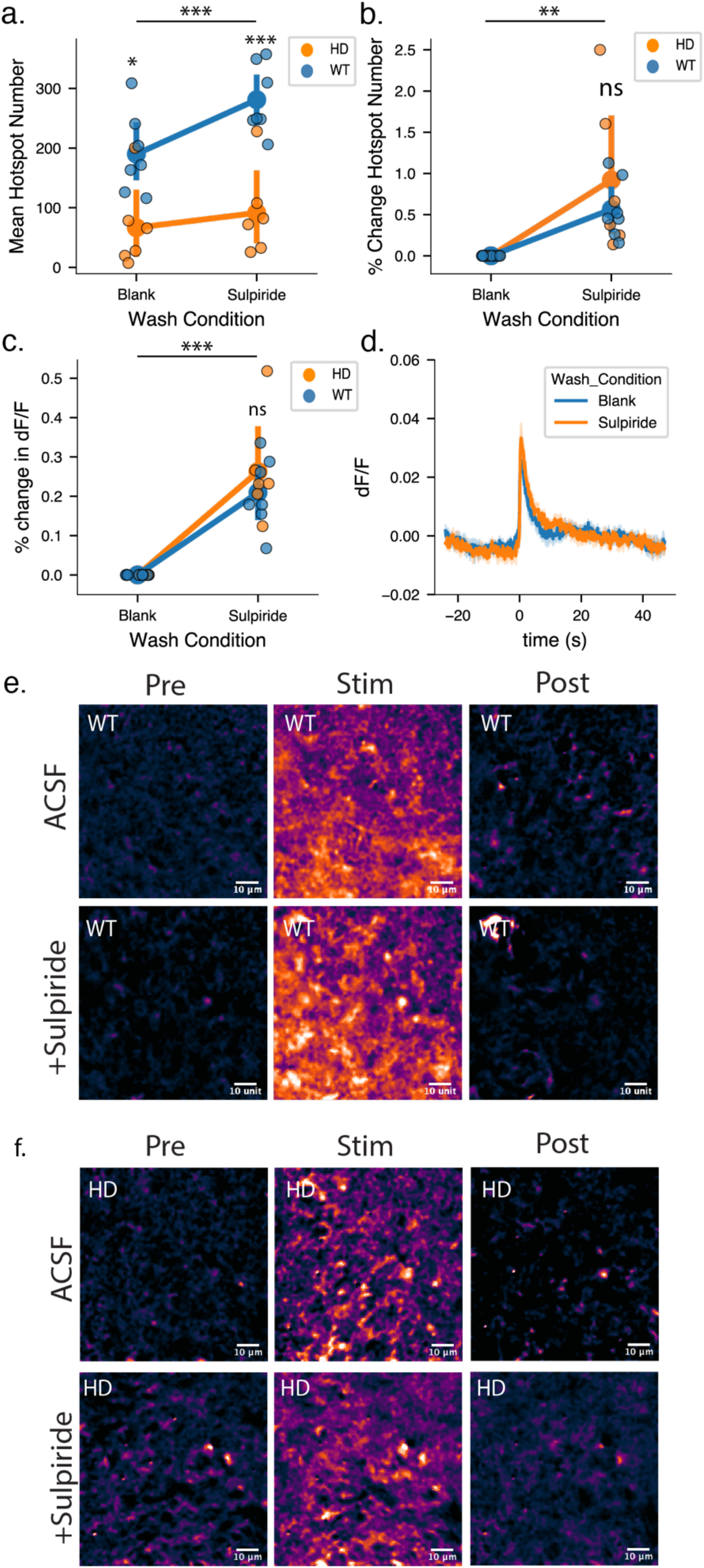
R6/2 HD mice at 12 weeks show decreased sensitivity to modulation of dopamine release via D2-autoreceptor antagonist Sulpiride. ***A,*** WT mice show a significant increase in the number of active dopamine hotspots in response to 0.3 mA stimulation after Sulpiride wash. In contrast, R6/2 mice show a significantly blunted increase in active dopamine hotspots. (WT N = 7 slices, 7 animals, HD N = 6 slices, 6 animals; mixed-ANOVA: disease state, p = 0.001; wash condition, <u>***</u> p < 0.0005; interaction, p = 0.001; paired t-test: * p = 0.009 HD/Blank compared to WT/Blank, ** p < 0.0005 HD/Sulpiride compared to WT/Sulpiride) ***B,*** Sulpiride wash results in comparable percent increase of dopamine hotspots in 12 week R6/2 HD and WT mice. (WT N = 7 slices, 7 animals, HD N = 6 slices, 6 animals; mixed-ANOVA: disease state, p = 0.074; wash condition, p = 0.003; interaction, p = 0.369; paired t-test: ns p = 0.412 HD/Sulpiride compared to WT/Sulpiride). ***C,*** Both R6/2 HD and WT slices show similar increase in percent increase in peak ΔF/F after Sulpiride wash (WT N = 7 slices, 7 animals, HD N = 6 slices, 6 animals; mixed-ANOVA: disease state, p = 0.411; wash condition, p < 0.0005; interaction, p = 0.411; paired t-test: ns p = 0.429 HD/Sulpiride compared to WT/Sulpiride). ***D,*** Dopamine release and reuptake traces from imaged from 12 wk HD mice. Solid lines denote the average taken from all slices and light shaded bands represent one standard deviation from average behavior. A 1 ms, 0.3 mA stimulation is delivered at time = 0s. ***E,*** Representative images of dopamine release imaged in 12 week WT mice before, during, and after stimulated dopamine release in the presence and absence of Sulpiride. *F,* Representative images of dopamine release imaged in 12 week HD mice before, during, and after stimulated dopamine release in the presence and absence of Sulpiride.

### Sulpiride promotes increased firing fidelity of ΔF/F hotspots in R6/2 HD and WT mice

The coverage of dopamine signaling across the striatum is influenced not only by hotspot number and peak ΔF/F, but also the fidelity of hotspot release. Here we term “hotspot release fidelity” as the ability of the same dopamine hotspot to fire upon repeated stimulations. To examine hotspot release fidelity, we utilized our ability to track individual dopamine hotspots across stimulations and recorded the number of stimulations out of three total that each hotspot was active (Fig. 6A). As such, hotspots that responded in all three stimulations were assigned a hotspot release fidelity of 3, while hotspots responsive in only one of three stimulations were assigned a hotspot release fidelity of 1. We then pooled all hotspots identified across 7 WT slices and 6 HD slices at 12 weeks and examined the distribution of hotspots across the three hotspot release fidelities before and after Sulpiride wash. As noted previously, Sulpiride can modulate dopamine release through increasing the number of dopamine hotspots or modulating the activity of existing hotspots (Fig. 5a, Fig. 5b). Therefore, we separated hotspots into those that are active before and after Sulpiride wash (“shared hotspots”) and those that emerge after Sulpiride wash (“added hotspots”).

**Figure 6.**
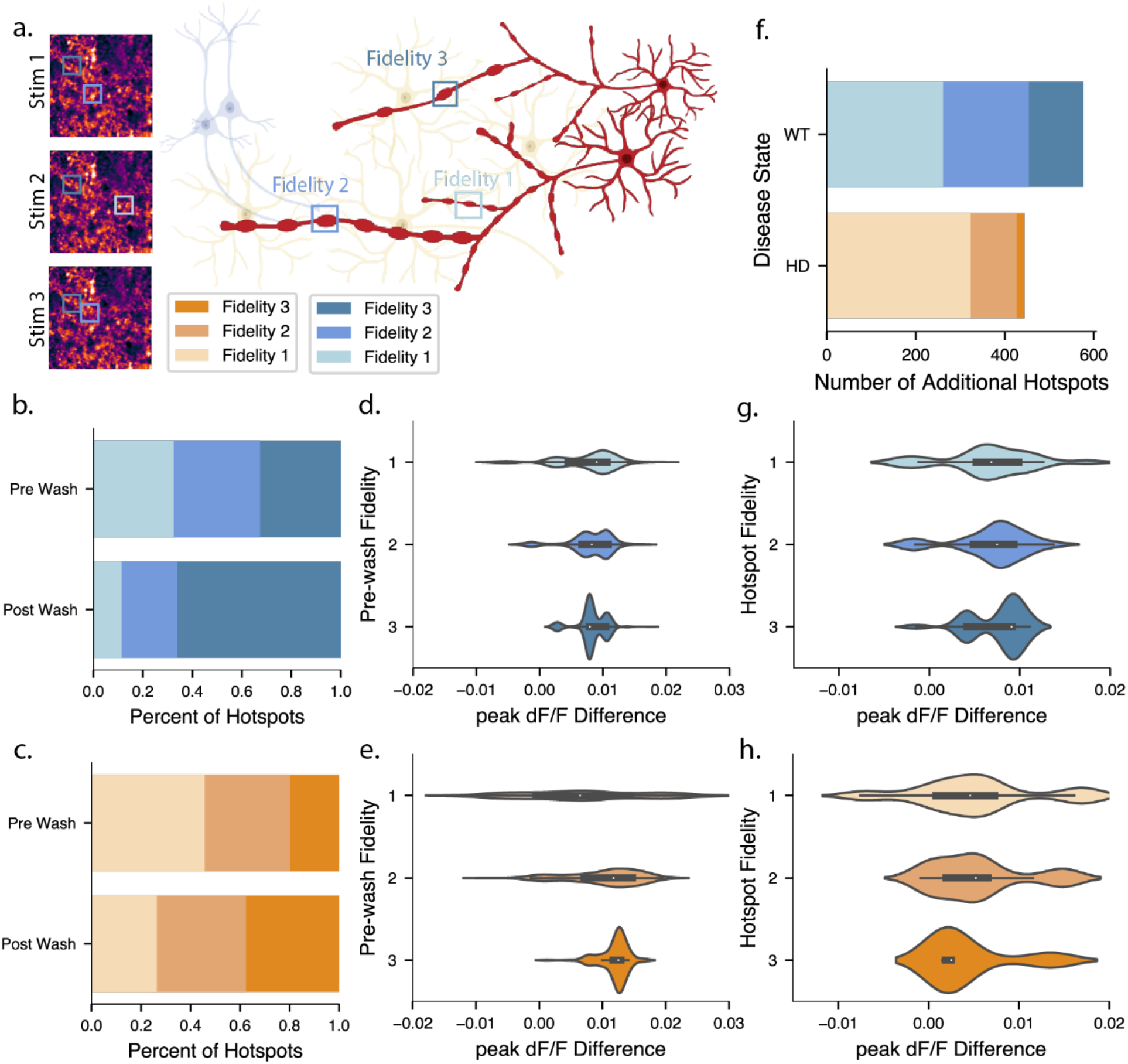
Sulpiride promotes increased firing fidelity of ΔF/F dopamine hotspots in both R6/2 HD and WT mice. ***A,*** Graphical overview of how individual dopamine hotspots can be tracked across stimulation replicates and assigned fidelity scores based on the number of stimulations that are active in. ***B,*** Stacked bar plot showing the distribution of shared dopamine hotspots active both before and after Sulpiride wash in WT 12 week mice (3836 dopamine hotspots total, pooled from 7 slices from 7 animals). Before Sulpiride wash 12 week WT dopamine hotspots are even distribution across fidelity scores (dark blue: fidelity 3, mid blue: fidelity 2, light blue: fidelity 1). After Sulpiride wash, fidelity 3 dopamine hotspots increase from making up 32.7% of all dopamine hotspots to 66.1% of all hotspots (pairwise tukey: <u>**</u> p = 0.002). This is paired with a decrease in fidelity 2 and fidelity 1 hotspots. ***C,*** Stacked bar plot showing the distribution of dopamine hotspots active both before and after Sulpiride wash in HD 12 week mice (1094 dopamine hotspots total, pooled from 6 slices from 6 animals). Before Sulpiride wash the majority of 12 week HD dopamine hotspots are fidelity 1 hotspots. (dark orange: fidelity 3, mid orange: fidelity 2, light orange: fidelity 1). Compared to fidelity 3 hotspots in WT slices, fidelity 3 hotspots in HD slices make up 13.0% less of the total dopamine hotspot population (pairwise tukey: <u>*</u> p = 0.035). After Sulpiride wash, 12 week HD slices do not show a significant increase in fidelity 3 dopamine hotspots (pairwise tukey: p = 0.308). ***D,*** Stacked violin plot showing the increase in dopamine hotspots mean peak ΔF/F after Sulpiride wash of WT dopamine hotspots active before and after Sulpiride wash. Values are sorted by the initial fidelity exhibited by the dopamine hotspot pre-Sulpiride wash. ***E,*** Stacked violin plot showing the increase in hotspots mean peak ΔF/F after Sulpiride wash of HD dopamine hotspots active before and after Sulpiride wash. Values are sorted by the initial fidelity exhibited by the dopamine hotspot pre-Sulpiride wash. ***F,*** Stacked bar plot showing the number of dopamine hotspots *added* by in HD and WT slices after Sulpiride wash. HD and WT Slices add a comparable number of dopamine hotspots after Sulpiride wash (pairwise tukey: p = 0.548). However, fidelity 3 hotspots make up a significantly higher percentage of added hotspots in WT slices compared to HD slices (pairwise tukey: <u>*</u> p = 0.004). ***G,*** Stacked violin plot showing the increase in hotspot mean peak ΔF/F after Sulpiride wash of WT dopamine hotspots *added* after Sulpiride wash compared to the average mean peak ΔF/F of all hotspots active before Sulpiride wash. Values are sorted by the fidelity exhibited by the dopamine hotspot after it appears following Sulpiride wash. ***H,*** Stacked violin plot showing the increase in hotspot mean peak ΔF/F after Sulpiride wash of HD dopamine hotspots *added* after Sulpiride wash compared to the average mean peak ΔF/F of all hotspots active before Sulpiride wash. Values are sorted by the fidelity exhibited by the dopamine hotspot after it appears following Sulpiride wash.

Before Sulpiride wash, WT 12 week mice show an even distribution of dopamine hotspots across fidelities. This distribution shifts following Sulpiride wash, resulting in an increase in the percentage of high release fidelity 3 hotspots from 31.0% of all dopamine hotspots to 66.6% of all hotspots (pairwise Tukey: <u>**</u> p = 0.0016) (Fig. 6b). In contrast, high release fidelity 3 hotspots represent 23.7% less of the of total dopamine hotspot population (pairwise Tukey: <u>*</u> p = 0.0137) in HD 12 week mice compared to WT 12 week mice. Furthermore, 12 week HD slices do not show a significant increase in fidelity 3 dopamine hotspots after Sulpiride wash (pairwise Tukey: p = 0.197) (Fig. 6c). Given that the number and identity of the shared hotspots is held constant before and after Sulpiride wash, increases in high release fidelity 3 hotspots in response to Sulpiride wash is driven by lower release fidelity hotspots transitioning into high release fidelity 3 hotspots. These findings are consistent with hypotheses that altered signaling through the D2-autoreceptor may alter voltage sensitivity of 4-amino-pyradine (4-AP) sensitive K^+1^ channels such as K_v_1.2 to shape the responsiveness of dopamine release (Fulton et al., 2011). We also examined whether a dopamine hotspot’s initial release fidelity in the absence of Sulpiride changed its modulation in peak dopamine ΔF/F following Sulpiride wash. Strikingly, we observed consistent modulation in dopamine hotspot peak ΔF/F regardless of initial release fidelity in both HD and WT slices (Fig. 6d, Fig. 6e), which suggests increases in fidelity are not a result of increased dopamine release leading to more consistent detection and that mechanisms leading to increase fidelity are separate from those increase peak ΔF/F.

We next examined the activity of hotspots *added* after Sulpiride wash. Interestingly, when controlling for the unique identity of dopamine hotspots, we found that HD and WT slices add comparable number of dopamine hotspots after Sulpiride wash (pairwise Tukey: p = 0.836) (Fig 6f). However, release fidelity 3 and release fidelity 2 hotspots make up 13.7% more and 10.7% more of added hotspots in WT slices in comparison to HD slices (pairwise Tukey: <u>*</u> p = 0.016, pairwise Tukey: <u>*</u> p = 0.035). Modulation of peak ΔF/F in added dopamine hotspots in comparison to the average peak ΔF/F of hotspots before Sulpiride wash was consistent regardless of the added hotspot’s release fidelity in both HD and slices and were not significantly different from that of shared dopamine hotspots (Fig. 6g, Fig. 6h).

These findings altogether suggest that the principal driver of decreased Sulpiride response in HD slices seen in Fig. 5a is changed dopamine hotspot release fidelity. Though HD slices add comparable numbers of unique dopamine hotspots after Sulpiride wash, reduced transition of dopamine hotspots to higher fidelity release combined with decreased addition of higher fidelity dopamine release hotspots ultimately results in fewer hotspots active during a given stimulation. Furthermore, while the fidelity of dopamine hotspots is significantly changed between HD and WT mice over the course of disease, changes in the peak dopamine ΔF/F of hotspots are comparatively mild even late in disease at 12 weeks. Together, these results suggest that altered dopamine release in HD is characterized by degeneration of dopamine release processes such that spatial coverage of dopamine release across the striatum is reduced. Though exposing late disease HD slices at 12 weeks to high extracellular calcium concentrations or Sulpiride indicates that additional dopamine hotspots can be engaged via molecular rescue, full rescue of dopamine signaling likely necessitates intervention at earlier HD timepoints.

### Blocking voltage gated K^+1^ channels with 4-AP and Sulpiride co-wash increases dopamine hotspot fidelity in WT mice while decreasing dopamine hotspot fidelity in HD mice

D2-autoreceptor action on the voltage gated K^+1^ channel K_v_1.2 plays a critical role in facilitating D2-autoreceptor mediated regulation of axonal dopamine release (Fulton et al., 2011). K_v_1.2 is the most abundant K_v_ subunit in the mammalian brain and blocking via 4-AP has been shown to counteract the ability of quinpirole to decrease FSCV detected dopamine overflow (Fulton et al., 2011). To assess whether the observed reduction in HD slice response to Sulpiride D2-autoreceptor antagonism at 12 week is driven by disruptions in D2-autoreceptor actions on K_v_1.2 channels or significant downregulation of D2-autoreceptors in the Striatum as a response to the dopamine depletion in late HD, we co-washed sulpiride and the broad spectrum K_v_1 channel family blocker 4-aminopyradine (4-AP) on slices to see if direct blockade of K_v_1.2 could further increase dopamine release from HD slices. We also increased the number of stimulation replicates from 3 to 10 to capture a wider view of how dopamine hotspots fidelity shifts with pharmacological action.

As anticipated, WT slices showed a progressive increase in number of dopamine hotspots following initial Sulpiride wash and subsequent Sulpiride and 4-AP co-wash. (Fig. 7a, Fig. 7b). This increase is in part facilitated through the promotion of lower release fidelity hotspots to higher release fidelity states, resulting in a progressive shift in the distribution of WT dopamine hotspots from skewing heavily towards low fidelity states to more even distributions following Sulpiride and 4-AP drug wash (Fig. 7d). Intriguingly, while HD slices show an increase in dopamine hotspots and high-fidelity dopamine hotspot after 10 µM Sulpiride wash, co-wash of Sulpiride and 4-AP decreases the number of dopamine hotspots and high-fidelity dopamine hotspots (Fig. 7a, Fig 7b). This decrease is not the result in decreased mean peak ΔF/F of HD dopamine hotspots compared to WT dopamine hotspots (Fig. 7c). Rather, decreases in dopamine hotspot number appears to be driven by an inability of 4-AP to promote lower release fidelity hotspots to higher release fidelity states in the HD striatum. (Fig. 7d).

**Figure 7.**
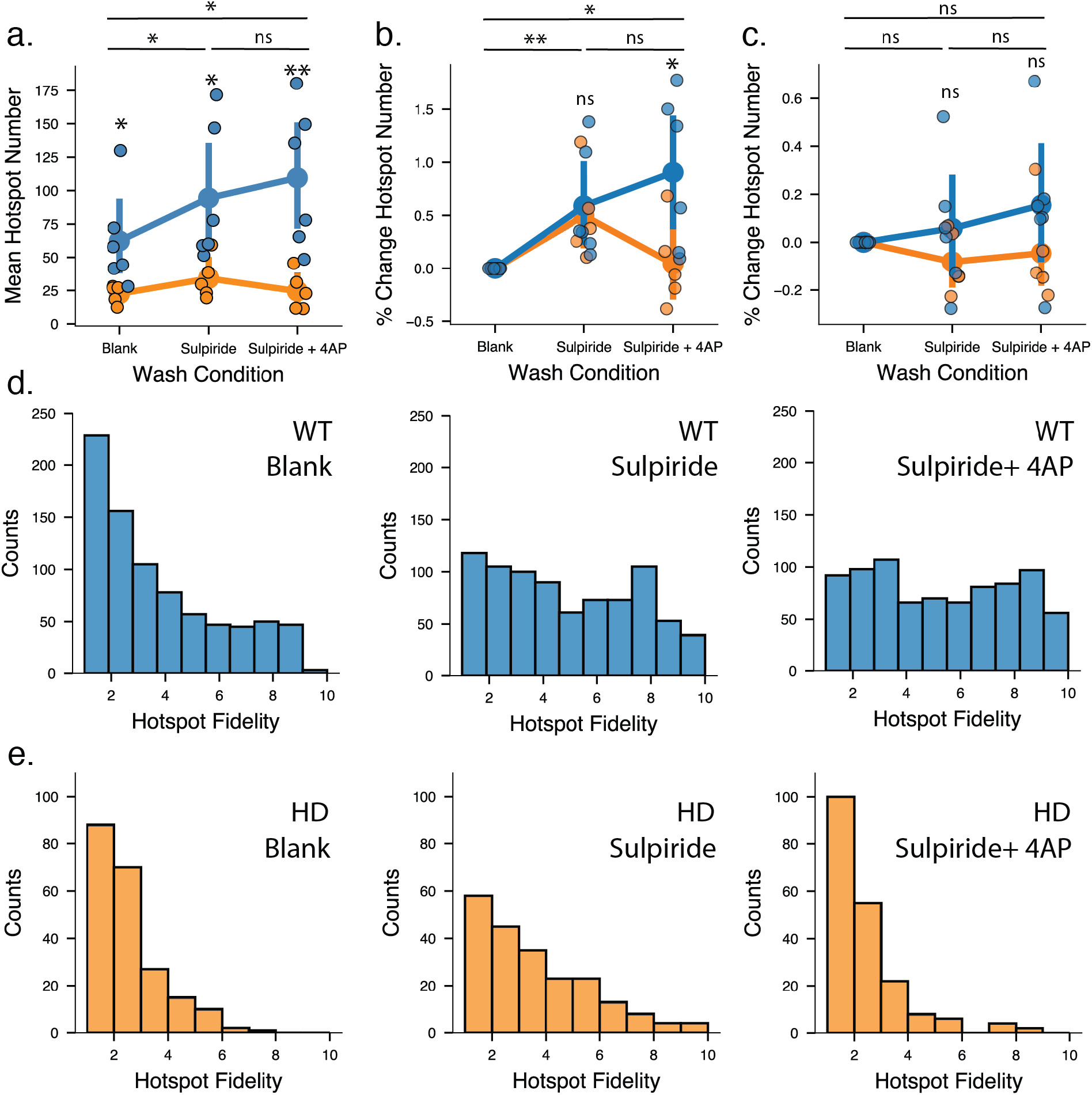
Sulpiride and 4-Aminopyridine (4-AP) co-wash increases dopamine hotspot fidelity in WT slices but decreases dopamine hotspot fidelity in HD slices. **A.** WT slices show a significant increase in the number of active dopamine hotspots over the course of progressive Sulpiride and 4-AP Wash. In contrast, HD mice show an increase in dopamine hotspots after sulpiride wash followed by a decrease in dopamine hotspots after 4-AP co-wash. (WT N = 6 slices, 6 animals HD N = 5 slices, 5 animals; mixed-ANOVA: disease state, *p = 0.014; wash condition, <u>**</u> p = 0.006; interaction, *p = 0.029; paired t-test: * p = 0.043 HD/Blank to WT/Blank, * p < 0.034 HD/Sulpiride to WT/Sulpiride, ** p < 0.001 HD/Sulpiride+4AP to WT/Sulpiride+4AP) **B.** HD and WT slices show comparable percent increase in dopamine hotspots after Sulpiride wash. However, HD slices show a striking departure in response after Sulpiride and 4-AP co-wash characterized by a decrease in dopamine hotspot number (mixed-ANOVA: disease state, p = 0.156; wash condition, <u>**</u> p = 0.003; interaction, *p = 0.020; paired t-test: p = 0.756 HD/Sulpiride to WT/Sulpiride, *p = 0.038 HD/Sulpiride+4AP to WT/Sulpiride+4AP) **C.** HD and WT slices show comparable percentx increase in mean peak dF/F after progressive Sulpiride and 4-AP wash (mixed-ANOVA: disease state, p = 0.264 wash condition, <u>**</u> p = 0.492; interaction, *p = 0.293; paired t-test: p = 0.299 HD/Sulpiride to WT/Sulpiride, p = 0.226 HD/Sulpiride+4AP to WT/Sulpiride+4AP) **D.** Histograms of pooled dopamine hotspots from all WT slices show that in blank ACSF dopamine hotspot distribution is skewed towards low release fidelity. Followed Sulpiride wash, dopamine hotspots increase in release fidelity, resulting in a more even distribution. This is further increased by Sulpiride and 4-AP co-wash. (permutation test on skew(post wash) – skew(pre-wash): statistic = −0.603 *** p < 0.0005 Blank/Sulpiride, statistic = - 0.130 * p = 0.045 Sulpiride + 4AP/Sulpiride) **E.** Histograms of pooled dopamine hotspots from all HD slices show that in blank ACSF dopamine hotspot distribution is skewed towards low release fidelity. Followed Sulpiride wash, dopamine hotspots increase in release fidelity, resulting in a more even distribution. However, this increase in release fidelity is lost after Sulpiride and 4-AP co-wash. (permutation test on skew(post wash) – skew(pre-wash): statistic = −0.364 p = 0.085 Blank/Sulpiride, statistic = 1.2 ** p ∼ 1.0 Sulpiride + 4AP/Sulpiride)

4-AP has also been documented to facilitate the opening of voltage gated Ca^+2^ channels to increase dopamine overflow outside of action on K_v_ channels (Wu et al., 2009; Fulton et al., 2011). However, the increased extracellular Ca^+2^ sensitivity we observe in HD slices at 12 weeks suggests that direction action on voltage gated Ca^+2^ channels should increase dopamine hotspot number and mean peak ΔF/F in HD slices. Therefore, 4-AP facilitated opening of voltage gated Ca^+2^ channels cannot account for the observed decrease in dopamine release in HD slices. Altogether, these findings point to a disease state in late HD where the ability of D2-autoreceptors to effectively regulate the release of axonal dopamine via K_v_1.2 is compromised. Rectifying this critical regulator of dopamine release would require direct and early targeting of these nigrostriatal dopaminergic neurons to restore proper dopamine signaling in the dorsal striatum.

## DISCUSSION

In this work we investigated spatial changes in dopamine release over the course of disease in R6/2 Huntington’s Disease model mice using nIRCat nanosensors. The synaptic-scale spatial resolution of these dopamine sensors enables identification of dopamine hotspots that are both sensitive to extracellular calcium concentration and D2R-autoreceptor antagonism. We show that progressive decreases in R6/2 HD dopamine release are driven by decreases in dopamine hotspot number, individual release site performance, and hotspot release fidelity. Early in disease, dopamine hotspots in HD slices show comparable extracellular calcium sensitivity and response to Sulpiride as WT mice. As disease progresses, the number of dopamine hotspots active in HD slices significantly decreases. Though increasing extracellular calcium concentration enables some increase in HD dopamine hotspot number by moving previously inactive dopamine hotspots into active states, increased calcium is not sufficient to restore dopamine hotspot number to WT levels. HD slices also demonstrate blunted response to Sulpiride antagonism of D2R-autoreceptors, which manifests primarily through decreased dopamine hotspot release fidelity. Interestingly, though the mean peak dopamine ΔF/F of hotspots in late disease HD slices is lower than that of WT slices, modulation of the mean peak dopamine ΔF/F of late HD hotspots via increase extracellular calcium concentration or to Sulpiride antagonism of D2R-autoreceptors is similar to WT dopamine hotspots. Altogether, these new spatial insights complement previous work exploring in the role of dopamine in Huntington’s Disease and build upon compelling evidence that molecular-level dopamine release mechanisms may be disrupted in late HD.

### Spatially-dependent dysregulation of dopamine dynamics in Huntington’s Disease

Disrupted dopamine transmission during Huntington’s Disease has been well documented in both HD patients and many genetic mouse models. In particular, the R6/2 mouse model has been noted to exhibit progressive decreases in dopamine release and basal tone using bulk dopamine measurement tools FSCV and microdialysis (Johnson et al., 2006, 2007; Ortiz et al., 2010, 2011; Callahan and Abercrombie, 2011). Herein with nIRCat, we find that 12 week old HD mice release only 23% of WT dopamine levels, in alignment with levels previously reported in existing R6/2 dopamine literature (Fig. S1c) (Johnson et al., 2006, 2007; Ortiz et al., 2010, 2011; Callahan and Abercrombie, 2011). Notably, our spatial insights from nIRCat imaging allow this late disease state to be interrogated at the level of release sites, revealing that decreases in overall dopamine release are primarily driven by a decrease in the number of dopamine hotspots rather than decreased individual hotspot performance. Computational modeling of phasic and tonic dopamine release has indicated that activation of D1-receptors and D2-receptors is complex and reliant on “spheres of influence” rising from each dopamine releasing terminal (Dreyer et al., 2010; Beyene et al., 2017). Our findings indicate that the sphere of influence of dopamine terminals in late HD is undermined not only through decreased dopamine release at individual hotspots, but also by decreased coverage across the dorsal lateral striatum due to loss of active hotspots. This decrease in dopamine coverage across the dorsal lateral striatum may underlie the altered dopamine signaling to dopamine receptors on direct pathway D1-MSNs, pathway D2-MSNs, and glutamatergic cortico-striatal glutamate terminals (Bamford, 2004; Kung et al., 2007; Cepeda et al., 2014; Koch et al., 2018; Koch and Raymond, 2019).

### Dopamine release in HD mice shows increased sensitivity to exogenous calcium concentration early in disease

We also investigate how extracellular calcium concentration affects dopamine release early and late in disease. FSCV measurements in R6/2 HD mice at 12 week show associated decreases in dopamine release cannot be fully attributed to decreases in striatal dopamine content alone (Johnson et al., 2006), leading to the hypothesis that calcium-dependent release machinery is disrupted in HD results in altered dopamine release. While previous work has reported that 12 week HD and WT mice show comparable changes in peak dopamine release concentration in response to increasing extracellular concentration, we find using nIRCat imaging that HD slices at 12 weeks show an increased sensitivity to high extracellular calcium concentration through the addition of dopamine hotspots. These findings point to the insights gained through nIRCat’s increased spatial resolution, as the reliance of previous dopamine detection methods on spatially averaging release from multiple dopamine release sites likely resulted in the comparable calcium modulation of dopamine hotspot performance (mean peak dF/F) between HD and WT slices to mask underlying changes spatial changes in dopamine hotspots mobilization. Altogether, these results suggest spatial changes in the ability of HD slices to recruit new dopamine hotspots at high extracellular concentrations may also play a role in altered dopamine release in late HD.

Human HD patients show biphasic disruption in dopamine release, with early dopamine excess believed to contribute to early chorea and late dopamine insufficiency to bradykinesia. This biphasic dynamic has been traditionally difficult to capture in mouse models including R6/2 (Cepeda and Levine, 2020). Nevertheless, striatal tyrosine hydroxylase (TH) activity in R6/2 mice has been shown to be biphasic, with elevated activity at 4 weeks and diminished activity at 12 weeks (Cepeda and Levine, 2020). To assess whether calcium-dependent dopamine release may be disrupted at a pre-symptomatic timepoint, we examined the extracellular calcium sensitivity of dopamine release at 4 weeks before motor changes. Stimulated dopamine release at this early 4-week time point has been previously uninvestigated in R6/2 mice, whereas our study explicitly queries dopamine modulation at this per-symptomatic timepoint. Interestingly, nIRCat imaging at 2 mM Ca^+2^ shows no difference in dopamine hotspot number or mean peak ΔF/F, indicating that stimulated dopamine release is not bi-phasically elevated in R6/2 mice at 4 weeks. However, 4-week HD hotspots do show increased peak ΔF/F at high extracellular calcium concentrations. This increased sensitivity of dopamine-releasing calcium machinery may be particularly notable during this early critical period between birth to P28 when dopamine is known to actively shape MSN excitability and could have important implications in describing the mechanism of subsequent dopamine dysregulation at later HD timepoints (Lieberman et al., 2018). Though murine HD models do not typically capture early choreic events, certain rat and non-human primate HD models have been reported to exhibit tetrabenazine responsive choreic movements and to be well suited to study early HD dynamics (Jahanshahi et al., 2010; Zeef et al., 2014; Chan et al., 2015). The non-genetically encoded nature of nIRCat nanosensor may be particularly advantageous to explore biphasic dopamine release in these non-murine tissues.

### D2R-autorceptor activity is disrupted in late Huntington’s Disease

D2-autoreceptors expressed on dopaminergic boutons play an active role in regulating dopamine release by initiating molecular events within the active boutons themselves. While expression and transcription of striatal D2 receptors is decreased in both HD patients and R6/2 mice, isolation of D2-autoreceptor behavior from dopamine receptors on neighboring cells remains challenging (Ariano et al., 2002; Vashishtha et al., 2013). By utilizing nIRCat’s to track individual dopamine hotspots and pharmacological compatibility, we are able to probe D2-autoreceptor action by via wash on of the D2R antagonist Sulpiride. In WT slices, Sulpiride antagonism of D2-autoreceptors shifts dopamine hotspots to higher release fidelities and promotes the activation of previously inactive high-fidelity dopamine hotspots. Notably, we find that high release fidelity dopamine hotspots comprise of 31% of all detected dopamine hotspots in WT slices. Our findings from nIRCat imaging stand in marked agreement with findings that only ∼20% of dopamine varicosities release dopamine and only ∼30% of dopamine varicosities are equipped with active zone-like machinery (Pereira et al., 2016; Liu et al., 2018). Conversely, HD slice show a weakened ability to convert dopamine hotspots into higher release fidelities. This ultimately results in fewer active dopamine hotspots in response to stimulation and decreased dopamine coverage across the dorsal lateral striatum.

These findings suggest possible disruption in the ability of D2-autoreceptors to work through K_v_1.2 channel pathways to shape the dopamine release or a striatal state in late HD where D2R dopamine pathways surrounding dopamine release have been downregulated such that further disinhibition does not dramatically increase dopamine release. We provide compelling evidence for disrupted K_v_1.2 channel activity, showing that selective blockage of K_v_1.2 channels with 4-AP combined with Sulpiride antagonism of D2-autoreceptors is unable to promote low fidelity dopamine hotspots to higher fidelity states. Conversely, in WT slices, we compellingly visualize that progressive wash of sulpiride and 4-AP shifts the distribution of dopamine hotspots from a population heavily skewed towards low fidelity hotspots to one of comparatively equal distribution between low and high hotspot fidelities.

We also find that dopamine hotpots in in HD and WT mice show similar changes in peak ΔF/F in response to Sulpiride. Interestingly, our Sulpiride-induced modulation of dopamine hotspot peak ΔF/F measured in the dorsal lateral striatum of R6/2 HD and WT mice is smaller than previously recorded modulations in the dorsal medial striatum of C57BL/6J mice (Beyene et al., 2019). This finding may indicate geographic differences in dopamine hotspot behavior across the striatum. We also observe that D2-autoreceptor response to Sulpiride changes in WT animals between 4 week and 12 weeks of age. While expression of dorsal lateral striatal D2-autoreceptors is known to increase and exhibit heightened sensitivity over the course of adolescence in rats, we find the opposite trend in WT R6/2 mice, with 4 week mice exhibiting lower response to Sulpiride in comparison to 12 week mice (Pitts et al., 2020). These differences may be a result of studying dopamine release in response to single stimulations, which contribute minimally to basal dopamine levels on D2Rs, rather than stimulation trains.

### Implications for potential therapeutic strategies for Huntington’s Disease Treatment

Huntington’s Disease is believed to manifest through the collective result of cell autonomous events and disruptions in synaptic signaling that compound into greater dysfunction (Cepeda and Levine, 2020). Global suppression of mutant huntingtin—even after symptom onset— has shown to facilitate molecular and behavior recovery in HD model mice (Yamamoto et al., 2000). Ongoing efforts to find effective treatments or cures for Huntington’s Disease have primarily focused on directly addressing the production of disease-causing mutant huntingtin protein at levels across translation, transcription, and the gene itself in striatal and cortical regions (Machida et al., 2006; McBride et al., 2011; Fink et al., 2016; Evers et al., 2018).

Coordinated targeting of mutant huntingtin in these compartments together has been shown to have a synergistic effect, strengthening the hypothesis that striatal cortico-striatal synapses play a critical role in HD pathogenesis (Wang et al., 2014). Our findings suggest that function of dopaminergic neurons that natively modulate this cortico-striatal signaling are a promising new therapeutic area that experience dysfunction during HD and may confer synergistic effects. In addition, our findings point towards the relevance of huntingtin directed or cell replacement therapies for repairing dopamine function, as the ability of dopamine receptor directed small molecule such as Sulpiride to modulate dopamine release to WT levels is diminished late in disease. Lastly, our results suggest that therapies may need to be delivered early in disease as aberrations in dopamine release are observed even before symptom onset.

## Acknowledgements

We are grateful for the technical assistance provided by Linda Wilbrecht and Kristen Delevich. We acknowledge support of a Burroughs Wellcome Fund Career Award at the Scientific Interface (CASI) (MPL), a Dreyfus foundation award (MPL), the Philomathia foundation (MPL), an NIH MIRA award R35GM128922 (MPL), an NIH R21 NIDA award 1R03DA052810 (MPL), an NSF CAREER award 2046159 (MPL), an NSF CBET award 1733575 (to MPL), a CZI imaging award (MPL), a Sloan Foundation Award (MPL), a USDA BBT EAGER award (MPL), a Moore Foundation Award (MPL), a DOE office of Science grant DE-SC0020366 (MPL), and an NSF Graduate Research Fellowship (S.J.Y.). MPL is a Chan Zuckerberg Biohub investigator, a Hellen Wills Neuroscience Institute Investigator, and an IGI Investigator.

## Notes

### Competing Interest Statement

The authors have declared no competing interest.

