## Supplemental Figures for "Synaptic scale dopamine disruption in Huntington’s Disease model mice imaged with near infrared catecholamine nanosensors"

SUPPLEMENTARY FIGURES

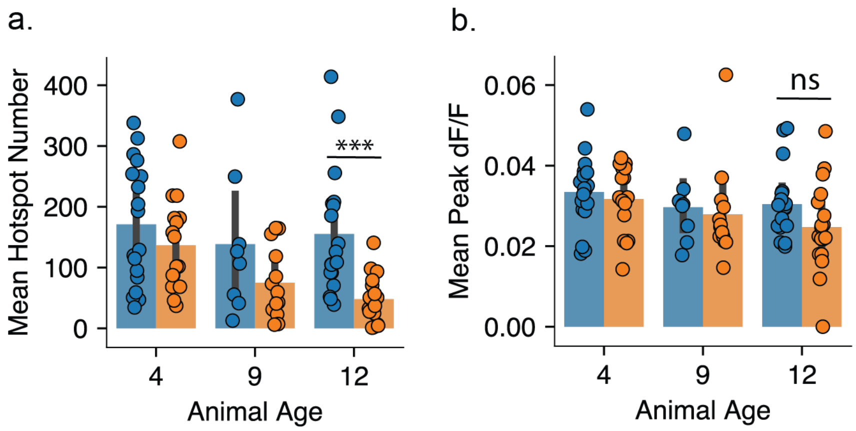


c.

| DA Detection Method | R6/2 HD DA release (% of WT DA release) | | |
|---|---|---|---|
| | **4 wk** | **9 wk** | **12 wk** |
| nIRCat Imaging (Hotspot Number x peak ΔF/F) | ~100% | ~84% | ~23% |
| Fast Scan Cyclic Voltammetry† | | ~60% | ~26% |
| Microdialysis†† | | ~43% | ~23% |

**Supplemental Fig 1.** ***A,*** R6/2 HD brain slices show progressively decreasing numbers of dopamine hotspots from 4 weeks through 9 and 12 weeks when stimulated at 0.1 mA while WT mice show no changes in dopamine hotspot number with age. (4 weeks WT N = 18 animals, HD N = 18 animals; 9 weeks WT N = 10 animals, HD N = 13 animals; 12 weeks WT N = 18 animals, HD N = 18 animals; ANOVA: disease state, $p = 0.106$; animal age, $p = 0.1013$ ; interaction, $p = 0.6419$; pairwise t-test: ns $p = 0.0921$ 12wk/HD compared to 12wk/WT, ns $p = 0.1969$ 9wk/HD compared to 9wk/WT, ns $p = 0.2627$ 4wk/HD compared to 4wk/WT). ***B,*** R6/2 HD brain slices show no change in average peak dopamine ΔF/F at 4 and 9 weeks when stimulated at 0.1 mA but show significant decrease late in disease at 12 weeks. (4 weeks WT N = 18 animals, HD N = 18 animals; 9 weeks WT N = 10 animals, HD N = 13 animals; 12 weeks WT N = 18 animals, HD N = 18 animals; ANOVA: disease state, $p <= 0.0005$; animal age, $p = 0.0309$ ; interaction, $p = 0.1982$; pairwise t-test: *** $p < 0.0005$ 12wk/HD compared to 12wk/WT, ns $p = 0.1969$ 9wk/HD compared to 9wk/WT, ns $p = 0.2627$ 4wk/HD compared to 4wk/WT). ***C,*** Amount of dopamine released in HD animals as a percentage of WT values was calculated via the equation (WT Hotspot Number x WT mean peak ΔF/F) / (HD Hotspot Number x HD mean peak ΔF/F). These values can be compared to values previously reported in literature. † (Johnson et al., 2006) † † (Callahan and Abercrombie, 2011)

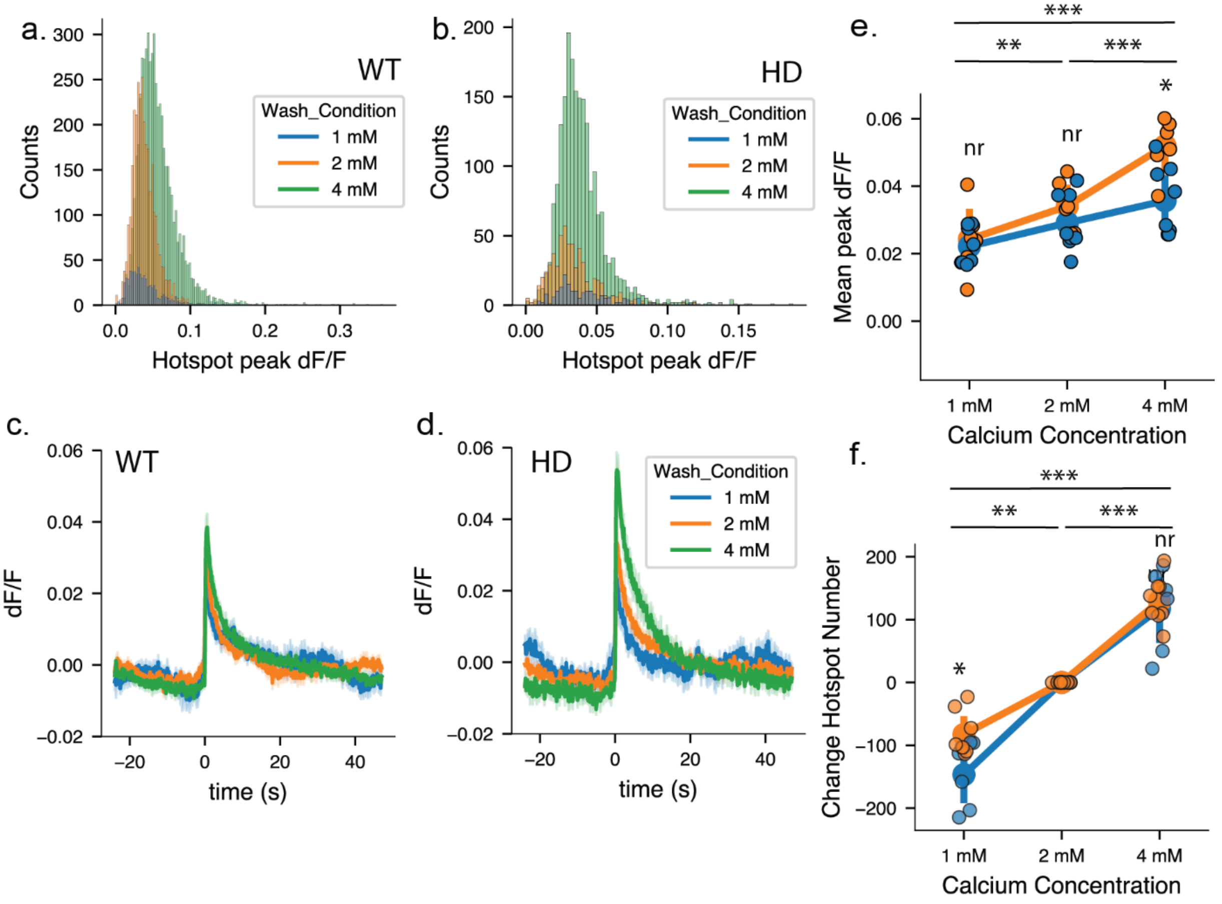


**Supplemental Fig 2.**

***A.*** Histograms of pooled dopamine hotspots from 4 week WT mic. Dark blue bars show hotspots active at 1 mM $Ca^{+2}$, orange bars hotspots active at 2 mM $Ca^{+2}$, and green bars hotspots active at 4 mM $Ca^{+2}$. ***B.*** Histograms of pooled dopamine hotspots from 4 week HD mic. ***C,*** Dopamine release and reuptake traces from imaged nIRCat-labeled brain slices for 4 week WT mice. Solid lines denote the average taken from all slices and light shaded bands represent one standard deviation from average behavior. A 1 ms, 0.3 mA stimulation is delivered at time = 0s ***D,*** Dopamine release and reuptake traces from imaged nIRCat-labeled brain slices for 4 week HD mice. ***E,*** The average mean peak ΔF/F values recorded in 4 week HD and WT slices at 1 mM $Ca^{+2}$, 2 mM $Ca^{+2}$, and 4 mM $Ca^{+2}$ ( mixed-ANOVA: disease state, *p = 0.1929; wash condition, *** p < 0.0005; interaction, p = 0.0093; pairwise t-test: ** p = 0.007 HD/1 mM$Ca^{+2}$ compared to WT/1 mM $Ca^{+2}$). ***F,*** Total change in hotspots number recorded in 4 week HD and WT slices at 1 mM $Ca^{+2}$, 2 mM $Ca^{+2}$, and 4 mM $Ca^{+2}$ ( mixed-ANOVA: disease state, *p = 0.0828; wash condition, *** p < 0.0005; interaction, p = 0.735; pairwise t-test: ** p = 0.0338 HD/4 mM$Ca^{+2}$ compared to WT/4 mM $Ca^{+2}$).

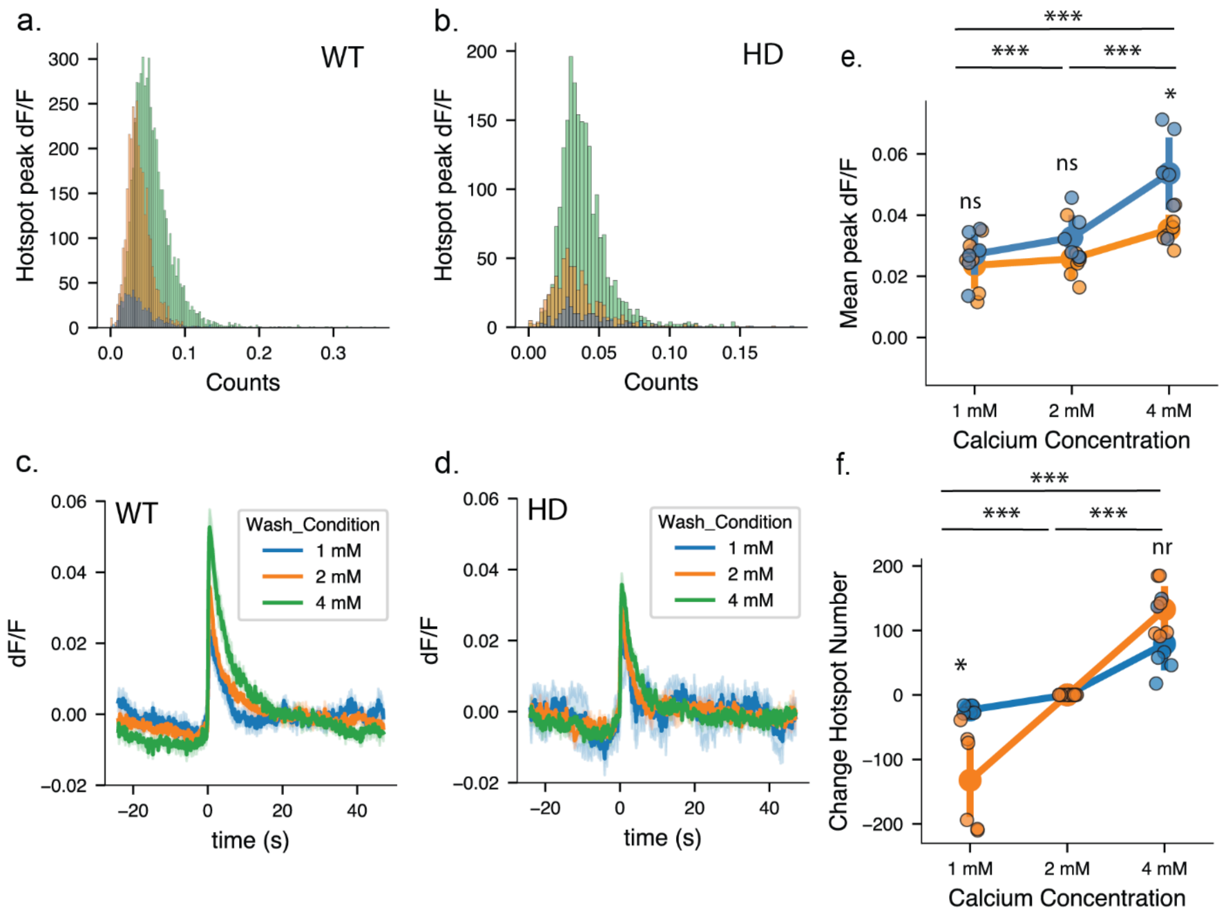


**Supplemental Fig 3.**
***A,*** Histograms of pooled dopamine hotspots from 12 week WT mic. Dark blue bars show hotspots active at 1 mM $Ca^{+2}$, orange bars hotspots active at 2 mM $Ca^{+2}$, and green bars hotspots active at 4 mM $Ca^{+2}$. ***B.*** Histograms of pooled dopamine hotspots from 12 week HD mic. ***C,*** Dopamine release and reuptake traces from imaged nIRCat-labeled brain slices for 4 week WT mice. Solid lines denote the average taken from all slices and light shaded bands represent one standard deviation from average behavior. A 1 ms, 0.3 mA stimulation is delivered at time = 0s ***D,*** Dopamine release and reuptake traces from imaged nIRCat-labeled brain slices for 4 week HD mice. ***E,*** The average mean peak ΔF/F values recorded in 4 week HD and WT slices at 1 mM $Ca^{+2}$, 2 mM $Ca^{+2}$, and 4 mM $Ca^{+2}$ ( mixed-ANOVA: disease state, *$p = 0.0471$; wash condition, *** $p < 0.0005$; interaction, $p = 0.0379$; pairwise t-test: * $p = 0.0164$ HD/4 mM$Ca^{+2}$ compared to WT/4 mM $Ca^{+2}$). ***F,*** Total change in hotspots number recorded in 12 week HD and WT slices at 1 mM $Ca^{+2}$, 2 mM $Ca^{+2}$, and 4 mM $Ca^{+2}$ ( mixed-ANOVA: disease state, $p = 0.268$; wash condition, *** $p < 0.0005$; interaction, $p < 0.0005$ ; pairwise t-test: * $p = 0.0074$ HD/1 mM$Ca^{+2}$ compared to WT/1 mM $Ca^{+2}$).

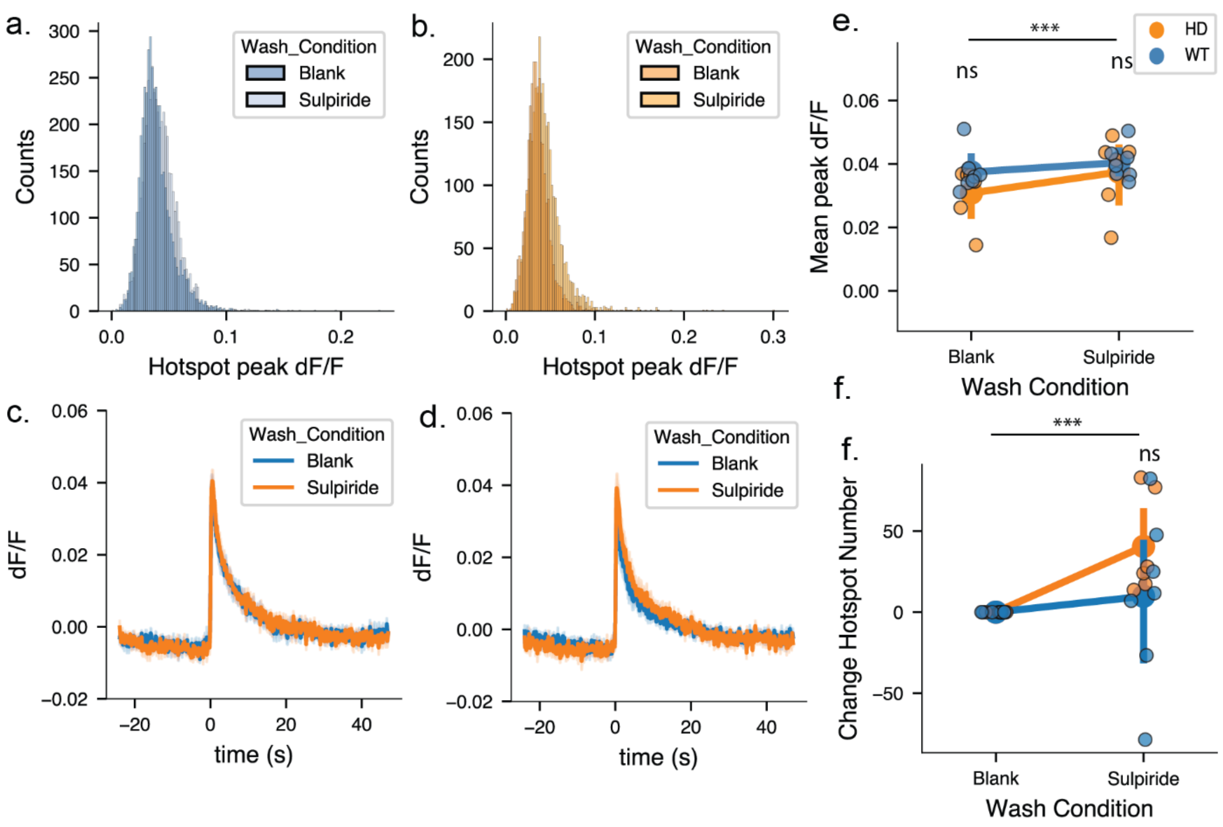


**Supplemental Fig 4.**

***A,*** Histograms of pooled dopamine hotspots from 4 week WT mic. Dark blue bars show hotspots active before Sulpiride wash, and light blue bars show hotspots active after Sulpiride wash. ***B.*** Histograms of pooled dopamine hotspots from 4 week HD mic. Dark orange bars show hotspots active before Sulpiride wash, and light orange bars show hotspots active after Sulpiride wash. ***C,*** Dopamine release and reuptake traces from imaged nIRCat-labeled brain slices for 4 week WT mice. Solid lines denote the average taken from all slices and light shaded bands represent one standard deviation from average behavior. A 1 ms, 0.3 mA stimulation is delivered at time = 0s ***D,*** Dopamine release and reuptake traces from imaged nIRCat-labeled brain slices for 4 week HD mice. ***E,*** The average mean peak ΔF/F values recorded in 4 week HD and WT slices before and after Sulpiride wash ( mixed-ANOVA: disease state, p = 0.310; wash condition, ** p = 0.002; interaction, p = 0.1466; pairwise t-test: ns p = 0.1676 HD/No Sulpiride compared to WT/No Sulpiride; ns p = 0.5927 HD/Sulpiride compared to WT/Sulpiride). ***F,*** Total change in hotspots number recorded in 4 week HD and WT slices after Sulpiride Wash ( mixed-ANOVA: disease state, p = 0.231; wash condition, ns p = 0.073; interaction, p = 0.231 ; pairwise t-test: ns p = 0.2170 HD/Sulpiride compared to WT/Sulpiride).

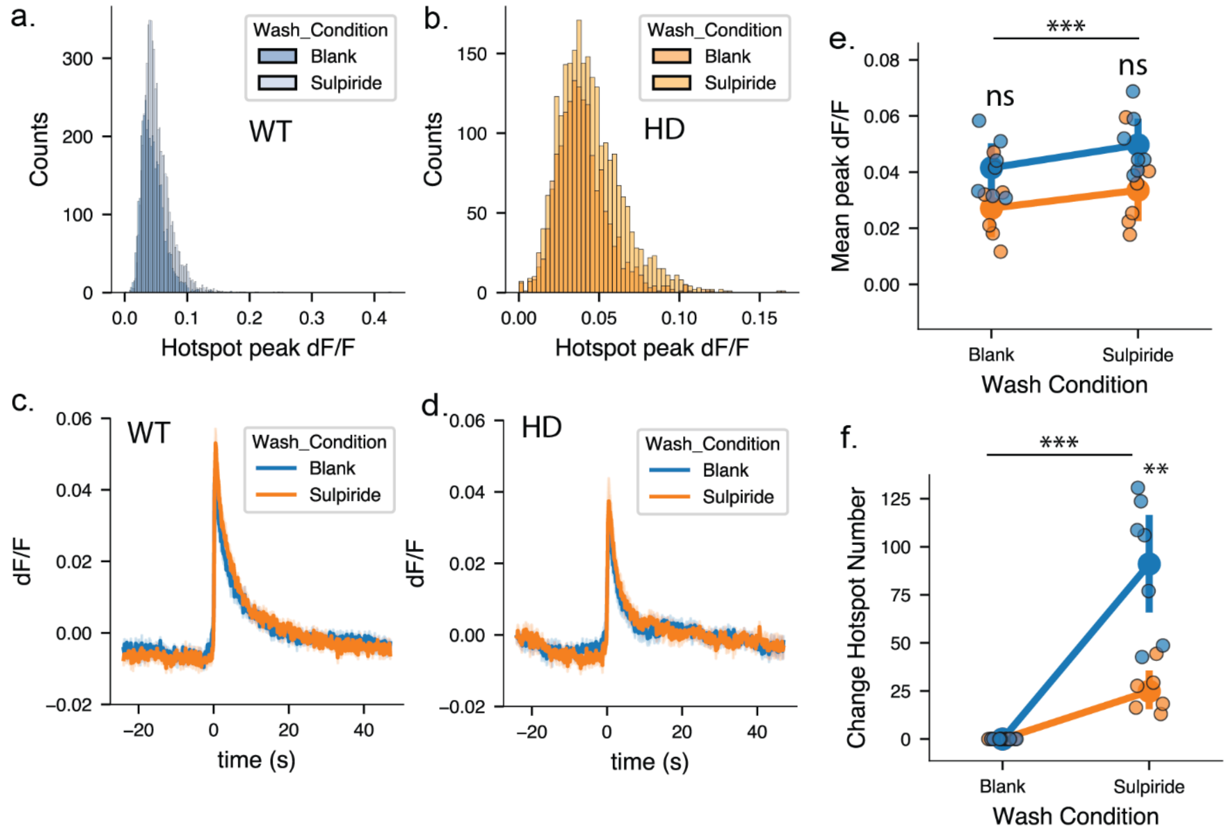


**Supplemental Fig 5.**

***A,*** Histograms of pooled dopamine hotspots from 12 week WT mic. Dark blue bars show hotspots active before Sulpiride wash, and light blue bars show hotspots active after Sulpiride wash. ***B.*** Histograms of pooled dopamine hotspots from 12 week HD mic. Dark orange bars show hotspots active before Sulpiride wash, and light orange bars show hotspots active after Sulpiride wash. ***C,*** Dopamine release and reuptake traces from imaged nIRCat-labeled brain slices for 4 week WT mice. Solid lines denote the average taken from all slices and light shaded bands represent one standard deviation from average behavior. A 1 ms, 0.3 mA stimulation is delivered at time = 0s ***D,*** Dopamine release and reuptake traces from imaged nIRCat-labeled brain slices for 4 week HD mice. ***E,*** The average mean peak $\Delta F/F$ values recorded in 4 week HD and WT slices before and after Sulpiride wash (WT N = 7 slices, 7 animals, HD N = 6 slices, 6 animals; mixed-ANOVA: disease state, $p = 0.0465$; wash condition, $p < 0.0005$; interaction, $p = 0.314$; paired t-test: * $p = 0.050$ HD/Blank compared to WT/Blank, ns $p = 0.059$ HD/Sulpiride compared to WT/Sulpiride). ***F,*** Total change in hotspots number recorded in 12 week HD and WT slices at 1 mM $Ca^{+2}$, 2 mM $Ca^{+2}$, and 4 mM $Ca^{+2}$ (WT N = 7 slices, 7 animals, HD N = 6 slices, 6 animals; mixed-ANOVA: disease state, $p = 0.001$; wash condition, $p < 0.0005$; interaction, $p = 0.001$; paired t-test: ** $p = 0.002$ HD/Sulpiride compared to WT/Sulpiride).
